# ML4SD: Leveraging Machine Learning and High-Throughput Search Algorithms for an Iterative Growth-Coupled Design Innovation

**DOI:** 10.64898/2026.09.11.750664

**Authors:** Alvaro Garantilla Becerra, Juan Nogales

**Affiliations:** Systems Biotechnology Group, Department of Systems Biology, Centro Nacional de Biotecnología, CSIC, Madrid, Spain; CNB DNA Biofoundry (CNBio), CSIC, Madrid, Spain; Interdisciplinary Platform for Sustainable Plastics towards a Circular Economy-Spanish National Research Council (SusPlast-CSIC), Madrid, Spain

## Abstract

Optimizing microbial biomanufacturing is required if renewable and waste carbon are to replace petrochemical routes at competitive titers, rates, and yields. Growth-coupled (GC) production supports that goal by linking target synthesis to biomass formation, so product formation is required for growth. Constructing knockout strains yielding GC production from a list of candidate genes is labor and time demanding. This results in few *in vivo* tested designs, which hampers standard machine-learning methods to learn GC patterns for specific bioprocesses. We therefore developed ML4SD, an active-learning Design–Build– Test–Learn (DBTL) cycle that trains ensembles on genome-scale metabolic model (GEM) scores of knockout designs, sampling the next designs from predicted model performance and error. That cycle generalizes only if the initial library is large and diverse, including suboptimal and non-viable designs; libraries restricted to minimal designs or Pareto-optimal knockouts were found to generate models overfitting. To meet those specific demands a novel strain design algorithm, gcSwarms, was developed and tested for a diverse set of bioprocesses. ML4SD was tested with an *in silico* case study converting lignin-derived 4-hydroxybenzoate to 6-caprolactam, the nylon-6 monomer. ML4SD results showed improvements of up to 164% on carbon yield, recovering a shared SHAP motif that redirects TCA flux through acetyl-CoA. Importantly it reaches that result using 2.5- to 7.1-fold fewer designs than a gcSwarms-only search, demonstrating the data efficiency of this method.

## Introduction

### Bioprocess optimization as the cornerstone of circular economy

Linear chemical extracts fossil carbon, converts it once, and discards the product, driving waste accumulation, greenhouse-gas emissions, and pollution with consequences for human populations and the biosphere. A circular bioeconomy instead keeps carbon in use by converting renewable biomass and waste streams into chemicals, materials, and fuels, reducing virgin petrochemical demand. Microbial bioprocesses sit at the center of that substitution: they can turn lignocellulose, industrial off-gases, municipal organics, and depolymerized plastics into marketable products rather than liabilities.^1^

However, a circular route is not necessarily a competitive one. For example, a bio-based product such as polyhydroxyalkanoate still sells at several times the price of fossil plastics. The costs of carbon substrates, pretreatment of recalcitrant biomass and downstream recovery often imply incurring more expenses than traditional chemical industry. Therefore, sustainable biomanufacturing means raising titers, rates, and yields on cheap, non-food carbon while keeping the whole process defensible under techno-economic and life-cycle assessment. Without that dual improvement in cost and sustainability, microbial conversion cannot displace petrochemistry at the scale a circular bioeconomy requires.^2^

Consequently, raising biomanufacturing performance is an active area of research. Classical campaigns still rely on empirical trial-and-error screening and on one-factor-at-a-time genetic or medium changes, which are labour-intensive and scale poorly as the design space grows. A promising alternative consist on the so called the iterative Design–Build–Test–Learn (DBTL) cycle. Basically, this methodology will start by specifying candidate genetic or bioprocess modifications (design). After that, those modifications will be implemented within a production host inhabiting a specific media (build). Once assayed (test), the measured outcomes are processed to extract patterns (learn) that will help to generate optimal modifications for the next round. Relative to sequential manual optimization, DBTL organizes that search so that combinatorial libraries can be reduced, bottlenecks identified systematically, and successive rounds executed on a shorter calendar. Biofoundries have adopted the cycle as one engineering option for biomanufacturing development, typically alongside chemical engineering and other process-development tools rather than as a replacement for them.^3,4^

Importantly, Machine learning (ML) changes what the learn step can do. Instead of inspecting results by hand, statistical and ML models map design factors to titers and return rules for the next build step, so fewer constructs need to be made and tested per round. In automated pipelines that coupling has already turned a second iteration around in weeks rather than in an open-ended screen. Those gains still leave DNA synthesis and poorly automated physical steps as delays, but they make DBTL a practical route from a laboratory conversion toward a process that is both cheaper and more defensible on sustainability grounds.^3^ In the context of biomanufacturing, the initial design step of that cycle still needs a predictive map of host metabolism for generating the first set of recommendations.

### GEM importance in metabolic engineering tasks

In that context, genome-scale metabolic models (GEMs) can provide a good approximation. GEMs have proved to be a powerful tools for simulating and analyzing the metabolic processes of organisms, effectively functioning as digital twins of their metabolism. These models can also be used for integrating various types of biological data, including genomic, transcriptomic, proteomic, and metabolomic information, to create comprehensive representations of metabolic networks. The integration of these diverse datasets allows for a more accurate depiction of an organism’s metabolic capabilities and responses to different environmental conditions or interventions^5^.

However, the predictions made from those platforms will always be incomplete, as there are still genomics sequences with poor or no annotation and enzymes with promiscuous interactions not considered that can make predictions sloppy^6^. Despite this incompleteness, the stoichiometric core of a GEM remains a usable scaffold for computational strain design (SD). Strain-design algorithms exploit gene–protein–reaction associations to propose gene interventions that redirect flux toward a target chemical, reducing reliance on trial-and-error screens.^7^

Those interventions are especially powerful when they enforce growth-coupled production, hardwiring target synthesis to biomass formation so that product formation is required for growth. Weak coupling requires a non-zero production rate at maximal growth; strong or obligatory coupling requires production at any feasible growth rate, which is intended to preserve synthesis even when cells grow suboptimally or experience local nutrient limitation.^8^ Because non-producing mutants would otherwise enjoy a selective advantage, coupling is argued to stabilize production during serial passaging and continuous cultivation.^8,9^ Once those gene interventions (knockouts or up/downregulations) are in place, adaptive laboratory evolution can act as a selection engine, driving regulatory and metabolic adjustments toward the computed growth–production phenotype.^8^

The identification of these intervention sets is typically approached as an optimization problem within a combinatorial design space, which is composed of the candidates for modification and the intervention type(s). Bilevel mixed-integer formulations such as OptKnock maximize an engineering objective in an outer problem while the inner problem represents growth. Alternative approaches such as minimal cut sets (MCS) identify the smallest deletion combinations that block alternative, uncoupled pathways.^7^ Furthermore, multi-objective genetic algorithms are employed to address this issue, with gcFront^10^ being a notable example. This algorithm, search Pareto fronts over growth rate, product synthesis, and coupling strength to speed up the search, outperforming other state-of-the-art methods. In any case, all those mixed-integer search methods are computationally expensive. So, to make the search less demanding, knockout-candidate computation typically prunes blocked and essential reactions, lumps unbranched pathways, or reuses database priors of previously computed designs.^11^

In any case, these stoichiometric tools remain limited in ways that bear directly on bioprocess robustness. Despite several GEM contextualization platforms exist, classical GEMs omit enzyme kinetics, protein allocation and regulation, and therefore overpredict cellular capability. However, thanks to recent advances in machine learning (ML), novel methodologies are starting to offer a feasible way to overcome those kinetic and regulatory gaps.^12^

### Machine Learning (ML) meets biotechnology

ML methods are beginning to assist bioprocessing tasks, and they do so most effectively when they complement genome-scale models rather than replace them. Implemented applications include campaigns that raise product titers: machine-learning recommenders have improved tryptophan production in Saccharomyces cerevisiae^13^ or isoprenol production in Pseudomonas putida.^14^ Other approach is to combine flux predictions from GEMs with a learner that ranks reactions associated with high-producing phenotypes, so the reconstruction maps feasible metabolism while ML highlights which parts of that map matter for production.^15^

Also, there exists a second class of applications that feeds experimental profiles back into the model. Metabolic-informed neural networks (MINNs) combine transcriptomic or proteomic measurements with a GEM so that predicted fluxes remain consistent with the reconstruction, allowing omics to inform the model without discarding its stoichiometric scaffold.^16^ Similarly, hybrid bioprocess modelling can embed a metabolic network inside a data-driven simulation of the fermentation, paving the way for a ML-assisted real-time control of bioprocesses.^17^ All those applications exploit the same fact: a ML element supplies the cellular context that stoichiometry alone cannot capture, while the GEM keeps the predictions biologically constrained.

However, despite the proliferation of multiple ML methods assisting metabolic engineering endeavors, these approaches have been mainly focused on protein and pathway engineering.^12,13,18^ Relatively little efforts have been dedicated to developing robust ML-based methods for metabolic optimization through gene deletion approaches.

To our knowledge, DeepGDel^11^ is the only published framework for this task, using modular deep learning to predict gene deletion statuses from sequential metabolite and gene representations. However, DeepGDel possesses notable limitations. Firstly, it does not explicitly account for the graph-based metabolic topologies, ignoring the connectivity between metabolites that provide a metabolic context for the bioprocess, which is widely used by strain design algorithms for finding deletions yielding GC phenotypes^11,19^ . Critically, DeepGDel predicts binary deletion statuses rather than the quantitative performance of a design—such as specific titers or yields—hampering its use for design optimization tasks or for *in vivo* validation.

The main challenge hampering the development of robust ML-based methods for metabolic optimization through gene deletion is the lack of SD data. As is widely known in the field, an ML model is only as good as the data provided to train it, so using enough high-quality data will be fundamental for any implementation to be successful. This arises as a problem when approaching an SD campaign, because the crafting of optimized strains is a time-consuming process which implies that only a few designs per bioprocess are available to learn from. For example, the implementation of DeepGDel, is constrained by a heavy reliance on extensive pre-existing databases for training, which drastically limits its scope to already computed or tested sets, not enabling *de novo* SD.

Recent efforts have been made in the development of high-throughput methods for both generation and screening of cell factories, with some promising applications derived from cell-free methods combined with CRISPR-dCas9 systems and microfluidics respectively^20–22^. Also, large language models (LLMs) are impacting the availability of high-quality SD data. An excellent example is D2Cell^23^ (Deep learning-aided Design of Cell factories), which is It is an artificial intelligence framework that uses LLMs to automatically extract metabolic engineering strategies from scientific literature focused on microbial strain design. Presently, it consists of a structured database of 29,006 cell factory designs across 1,210 unique products and 751 organisms, being the largest dataset of experimentally tested SD data. However, those techniques are still missing to fulfill the need of those data-hungry algorithms, and so the focus is on designing ML methods that can learn with less input data.

In this sense active-learning (AL) approaches have much to tell. This methodology, makes use of machine learning algorithms to determine the next set of experiments to be carried out while studying a given problem^24^. This has proved to be more data efficient in comparison with conventional ML approaches, which has led to some applications of them in bioproduction optimization^13,25^.

Also, as previously mentioned, SD can be seen as an optimization problem where our aim is to find a set of interventions yielding a best performance metric. Under this view, it is possible to formulate continuous learning cyclic frameworks capable of automating strain design and subsequently accelerating cell factories, following already developed technology. However, the development of such frameworks has not been fully approached and remains a major pursuit in bioengineering^4^.

Approaches like DeepGDel require high computational demands for retraining^26^, limiting its efficiency as a dynamic recommendation tool for continuous learning. Therefore, for tackling SD optimization, we propose an alternative approach consisting of a GEM-founded DBTL cycle specifically aimed at improving ML-guided gene knockout recommendations for growth-coupled strain design (**Figure 1**).

**Figure 1.**
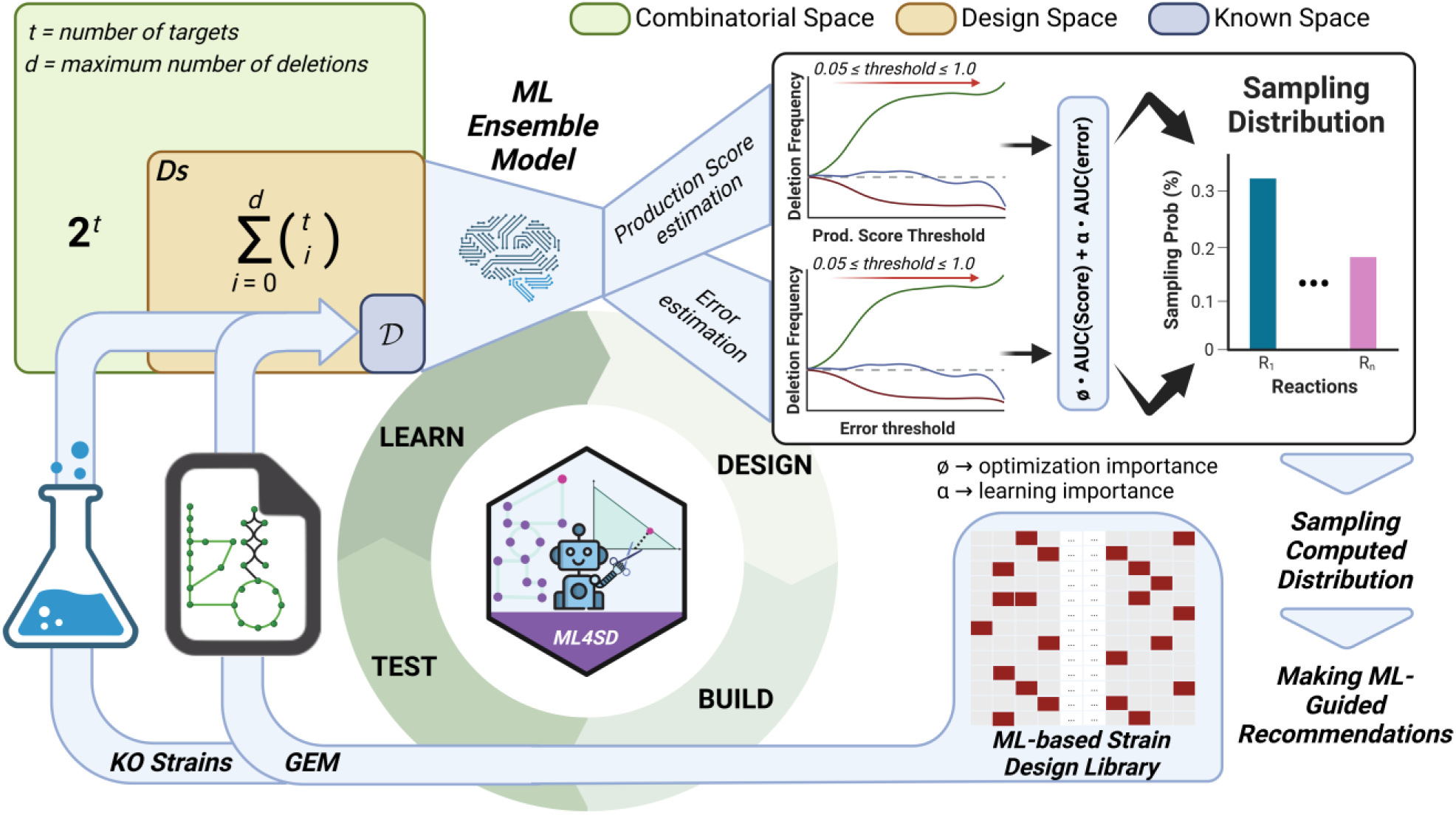
Proposed DBTL cycle for Strain Design through Machine Learning integration (ML4SD). Schematic representation of the ML4SD framework for ML-guided strain design optimization. The framework integrates an active-learning approach within a DBTL cycle to iteratively refine strain modifications coupling production to growth. Initial strain designs are generated using SD algorithms over high-quality GEMs, followed by validation through in silico or in vivo phenotyping. Ensemble Machine-Learning (ML) models, trained using *AutoSklearn*, predict performance scores and inform the next cycle’s modifications. Resultant recommendations are based on an adaptive sampling distribution computed from individual deletion contributions to ensemble error and/or scores prediction, enabling for explorative or exploitative modes of ML4SD respectively.

## Results

### 1 Towards a ML-guided active-learning framework for strain-designing empowered by GEMs

We build upon previous ML works^27,28^ and developed an active-learning (AL) approach for strain designing, which we called Machine Learning for Strain Design (ML4SD). In the absence of experimental data for a given bioprocess (e.g., a selected carbon source as input being bio transformed into a target metabolite as output), ML4SD leverages GEMs as generator of high-quality, high-consistent, and time-efficient *synthetic* data for generating ML models. ML4SD uses as a starting point an initial Strain-Design (SD) library of designs optimizing the target bioprocess which is denoted as D (known space,**Figure 1**). This library is generated using a reduced GEM, which helps identify a set of candidate reactions that guide SD algorithms to generate the Knockout-based designs (MM section 2.2). The designs in the library are represented as binary vectors of length equal to the number of different candidate reactions in the initial library (*t*). The maximum number of deletions, encoded as ones, is customizable and it can be set by the user (*d*):

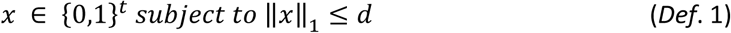

Considering this data structure, the total number of potential different designs (e.g., design space (*D_S_*)), is given by (2).

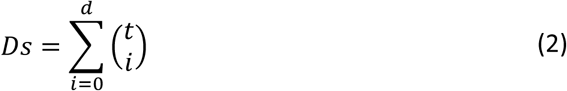

To evaluate the feasibility of the designs, we can either, apply an *in-silico* method using the complete GEM, or adopt an *in vivo* approach in a high-throughput phenotyping scenario (test step). Those scored designs constitute D (subset of *D_S_*), where different ML models are trained on (learn step).

For model ensembling, ML4SD employs *AutoSklearn*^29^ (https://github.com/automl/auto-sklearn) a Python package capable of training different types of ML models while optimizing their hyperparameters in a parallel and computationally efficient manner. In our case, each individual model is trained to associate a production metric with each design within D. Finally, among all trained models, the most accurate ones are selected and combined to form a final Ensemble Model that best fits the provided data^29^. It is important to note that the user can fine tune different aspects of both the learning process of the single ML model and construction of the final ensemble according to specific needs. For the specific setup we use in this study, including the cross-validation methodology, please refer to section 3 of materials and methods.

Using the predictions generated by the ensemble model, the individual contribution of each reaction, in terms of production score *(y*) and accuracy measured as absolute error (*AE(y)*), is quantified. These two values are further used to compute a sampling distribution as part of the design step (**Figure 1**). Specifically, within each cycle of ML4SD, the final sampling distribution (*S(*D)) is obtained by merging two individual distributions: one derived from *y* and the other from *AE(y)* according to (3).

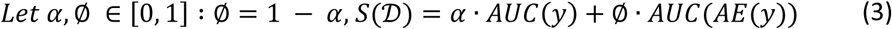

With the first term of the equation, we account for reaction deletions with high probability of optimizing the production score, while the second term provides information about deletions that results in low prediction accuracy. This enables us to guide subsequent ML4SD-fuelled DBTL cycles towards either an exploitative or an explorative goal, depending on the weight assigned to the learning or product optimization parameters (*α* or ∅ respectively). Guided by *S(*D), new ML-guided recommendations (additional KO designs) can thus be suggested for the next cycle contributing to a more comprehensive exploration of the design space while identifying improved designs. Importantly, these superior designs can be further validated either *in silico* using the complete GEM and/or *in vivo* through experimental implementation in the lab. This enable the integration of the GEM-based knowledge with experimental data, as envisioned in recent work^30^. These initial ML-models can further be fine-tuned with experimental data from promising strain design implementations (Test, Learn steps, **Figure 1**).

Taking all this into consideration, ML4SD could be, in principle, applied alongside others computational tools in biofactories using high-throughput phenotyping techniques to significantly accelerate the design of strains that overproduce metabolites of interest^4,31,32,15^. In the present work, we applied two different SD algorithms (gcFront^10^ and gcSwarms, discussed in following sections) to generate D for multiple bioprocess scenarios. With the resultant libraries, ML4SD approach was applied towards optimization of a biotechnologically relevant bioprocess. In the following, we describe in detail the different steps within ML4SD. All the code belonging ML4SD is available to any user by a free-access GitHub repository (https://github.com/extrevaro/ML4SD).

### 2 Exploring the feasible GC production space in *P. putida* to address the impact of bioprocess diversity on ML4SD

Before implementing ML4SD workflow we needed to know that ML models generated with the proposed methodology can perform well across a diverse metabolite library. Meeting this criterion will ensure the robustness and applicability range of the method. Considering that, we used the chemical space of *Pseudomonas putida KT2440* (*P. putida*) as a study case. *P. putida* is an ideal chassis given its carbon source utilization range and broad metabolic capacities^34^. Importantly, this organism has a high-quality GEM available, iJN1462. This model fulfills the requirements to be considered as a digital twin: comprising 1,462 gene products, 2,929 reactions, and 2,155 metabolites, with most reactions supported by primary literature, a MEMOTE consistency score of 97%, and experimental accuracies of 79% and 84% for carbon- and nitrogen-source growth screens.

It is important to note that we could generate fingerprints for 80% of all iJN1462 metabolites for further generating the embedding (MM section 1). As suggested in recent studies, we use the average fraction of 10^th^ nearest neighbors in the original high-dimensional data that are preserved as 10^th^ nearest neighbors in the embedding (*KNN*) as an indicator of the local data structure preservation^35^. Our chemical space has a *KNN* of 0.53 indicating that, as average, each metabolite shares more than a half of its fingerprint neighbors.

Considering the previous results, we can conclude that our generated chemical space correctly captures the chemical diversity within *KT* metabolism.

Substrates and products were selected to generate the different bioprocesses. The first ones were manually selected from the embedding, considering their presence as byproduct of other industries^36,37^ (molasses, lysates, lignocellulosic material…) and their chemical diversity. Metabolites classified as substrates were those in which the used GEM could grow on a minimal media using them as the unique carbon source (**Figure 2A**). For assesing their diversity, chemical dissimilarities were measured according to their Euclidean distances in the generated chemical space. Considering this, of 152 considered carbon sources in *KT*, 13 were selected (**Figure 2B**). Their median pairwise distance is 27% higher in comparison with all carbon sources (**Supplementary Figure 2**), therefore representing a diverse chemical set. On the other hand, 16 products were selected based only on their biotechnological potential on the selected chassis^38–40^.

**Figure 2.**
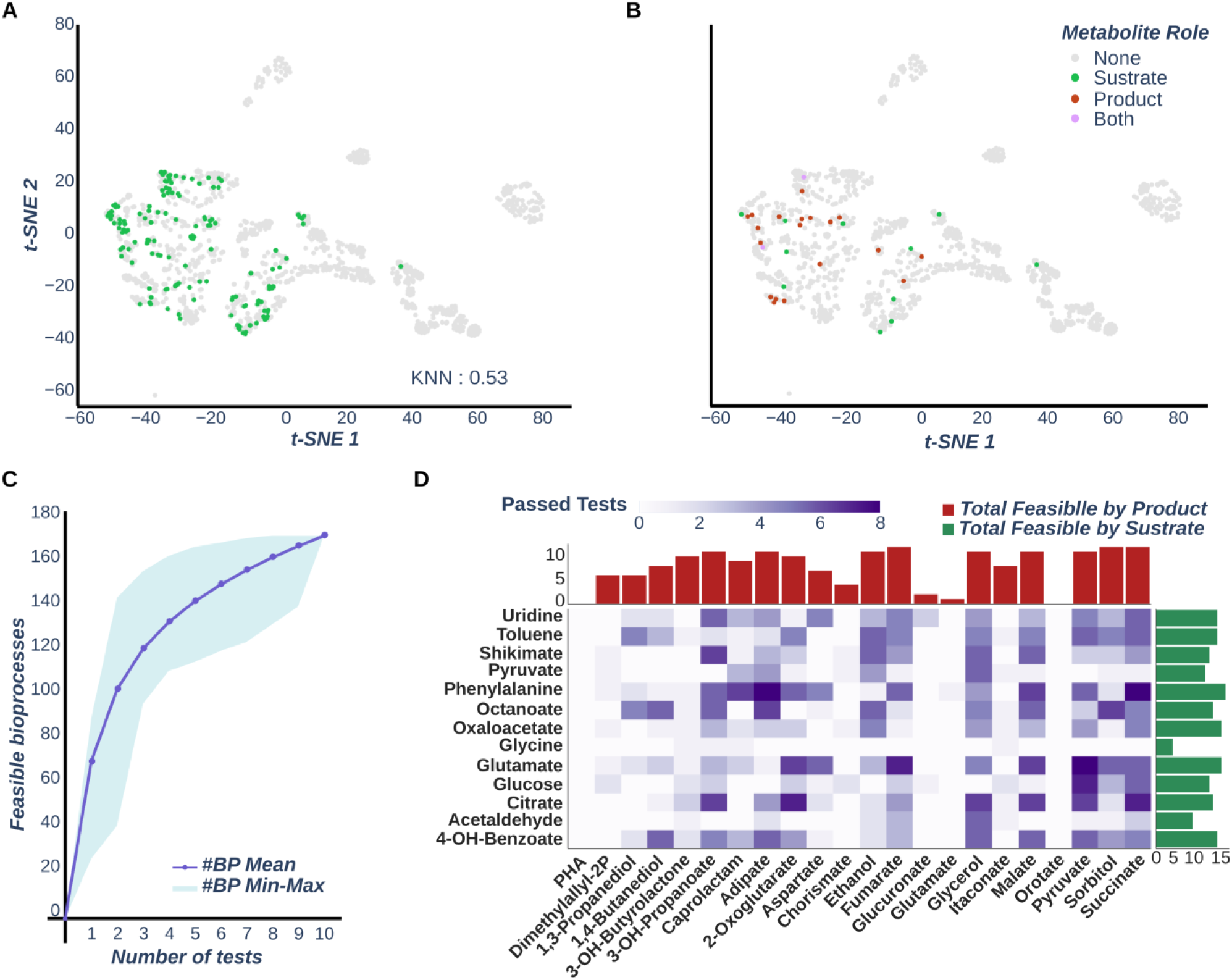
Coupling feasibility analysis over iJN1463 chemical space. **(A)** t-SNE representation of all *iJN1463* metabolites according to their fingerprints generated through *RDKit* python package.Metabolites identified as potential substrates are colored in green. (**B**) Same t-SNE representation but highlighting metabolites selected in this work for bioprocess generation, using the color code indicated in the legend. (**C**) Plot recapitulating the number of feasible bioprocesses relative to the number of MCS tests performed in the SD pipeline. Margins denote the range between minimum and maximum number of feasible processes within a given number of tests made, while the dotted lines represent the average across all MCS test runs combinations. (**D**) Heatmap showing the number of MCS tests passed by each bioprocess, with each row and column indicating its substrate and product respectively. Histograms are shown for substrates (green) and products (red), indicating the number of feasible bioprocesses for each metabolite.

With such a selection, we combinatorically generate a total of 284 different bioprocesses. As SD pipelines are known to be time-consuming and computationally expensive tasks, we filter bioprocesses following a minimal cut set (MCS) approach adapted from a previous study (see MM section 2.3). This test is applied up to 10 times (**Figure 2C, D**) for each selected bioprocess, using a different solver seed for each execution, which leads to a different exploration of the search space^41^.

The test results tell us that all substrates except glycine enabled at least one GC feasible bioprocess, with phenylalanine showing the highest number (17/22), closely followed by glutamate and ocaloacetate (16/22) and other aromatic compounds such as 4-hydroxybenzoate and toluene (15/22). Regarding products, 2 of them have no GC feasible bioprocesses (PHA and orotate), while TCA intermediates such as fumarate and succinate, along with the sugar sorbitol, stand as the ones with more (12/13) (**Figure 2D**). In summary, a total of 173 bioprocesses are considered feasible for coupling (61% of total), significantly reducing the number of tasks while maintaining chemical diversity of the metabolites involved. This amount of coupled bioprocess is within the range of a previous benchmark study when all metabolite outflows contained in the GEM are open, making our results on line with literature^41^. Therefore, we consider that the resultant battery of feasible *P.putida* bioprocesses represents a diverse enough set for testing the ML4SD computational framework.

### 3 Learning strain designs patterns by combining ensemble models and high-quality GEMs proves to be a challenge for state-of-the-art algorithms

Our framework starting point is the data from D, which is obtained with the aid of an SD algorithm applied over a high-quality GEM that serves as a digital twin of the chassis metabolism. Thus, it is vital that this initial data represents *D_S_* as best as possible. Otherwise, the resulting model will not properly capture the impact of deletion combinations over the phenotype, potentially propagating this bias through all DBTL rounds. An idea could be to generate comprehensive databases on deletion strategies for training the ML models. Recent efforts have been made in that direction, including novel algorithms integrating the data to alleviate the need of computational resources that is usually needed for SD tasks^42,43^. However, these approaches, though promising, are still in their infancy and have serious limitations. Importantly, the mentioned database MetNetComp (https://metnetcomp.github.io/database1/indexFiles/index.html) only contains designs coupling the production of metabolite from the default carbon source of the given GEM. We consider that the bioprocesses to optimize should only be limited by an expert consideration on its biotechnological relevance, not by preexisting deletion strategies data. This is why we have decided to use an SD method, enabling us to dynamically compute bioprocess-specific deletion strategies.

In this scenario, it is important that the strategy search performed by the SD method perform relatively fast while generating diverse data. Therefore, our first choice for this task was gcFront. This algorithm is designed to explore deletion strategies that simultaneously maximize cell growth, product synthesis and coupling strength (6) through three-level optimization. Making use of a genetic algorithm, gcFront can efficiently generate numerous optimal and sub-optimal alternative designs on the Pareto surface in a single run. In comparison with other SD algorithms, gcFront outperforms them in finding GC designs faster, making it a desirable start point^10^. Moreover, we are familiarized with the algorithm, successfully implementing it for experimentally addressing several biotechnological tasks^44,45^. Consequently, we execute our in-house SD pipeline (**Supplementary Figure 3**) using gcFront as SD algorithm over the different bioprocesses identified as feasible by our previous check, following the setup described in MM (Section 2.1-4). For those, we identified deletion strategies for 63 bioprocesses (50.4%), indicating that some of them were more challenging to address within our imposed computation time. Those libraries are composed of 20 to 349 viable designs (**Figure 4B**), which is a pretty small number in comparison with DS ^(^2^)^. For each of the generated libraries, we generate a bioprocess-specific ensemble model.

As previously said, for generating our ensemble models we have use the auto-sklearn python package. This module was designed to automate machine learning workflows, making them accessible to non-experts and improving their efficiency and robustness^29^. They optimize model selection by classifying the provided dataset according to its feature similarity and using historical performance data to provide optimal initial model configurations (meta-learning). For hyperparameter optimization of individual models, Bayesian optimization is used, while the ensemble construction is made from a weighted combination of all evaluated models during this search. This methodology increases the ensemble robustness by avoiding reliance on a single hyperparameter configuration. A basic scheme of the auto-sklearn ensemble construction can be seen in **Figure 3**. Overall, the mentioned characteristics make this platform a good candidate to deploy reliable machine learning models effectively.

**Figure 3.**
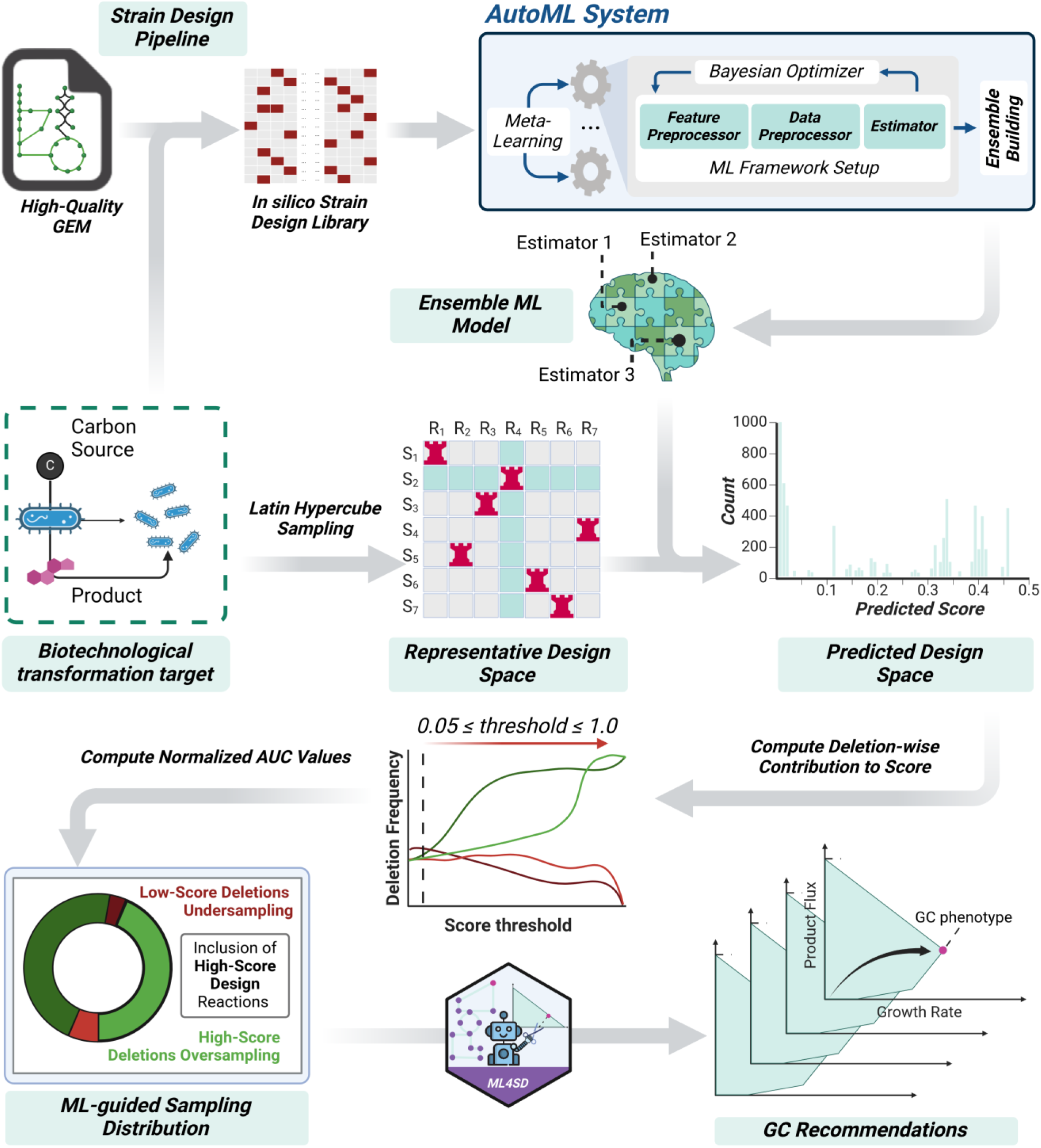
Schematic representation of ensemble model construction and the ML-based recommendation approach. *AutoSklearn* is used for ensemble generation. It leverages meta-learning for model selection, Bayesian optimization for hyperparameter tuning, and weighted ensemble construction for improved robustness. The resultant ensemble model is trained with SD libraries and used to generate SD recommendations. To this end, *LHS* is applied to generate a representative subset of possible strain designs within the design space (see MM section 4.2). The trained ML model evaluates these designs, identifying key deletions contributing to a performance or error score. Finally, by analyzing the frequency of each deletion across different score thresholds, an area under the curve (AUC) approach is used to derive a probability distribution guiding recommendation suggestion.

To train this ensemble, the libraries generated with gcFront were one hot encoded into a binary array of length *t* using the reaction candidate list generated in our *in-house* SD pipeline to fit the previously defined design data structure (*Def*. 1). Also, carbon yield (*CY)* is computed for each design according to ^(4^ and selected as the target variable to learn. For further details on ensemble model training setup, please refer to **Supplementary Figure 5** and section 3 of MM.

Once the models have been generated, we measured their predictive power through R^2^ of train and test sets. Considering the first, the models show a median R^2^ of 0.995 with little dispersion (**Supplementary Figure 5E**). However, despite the R^2^ values for the test set also indicates good predictive power (median of 0.952) of the model, their distribution varies significantly (**Supplementary Figure 5F**), having a median absolute deviation (MAD) 12 times higher. These results indicate bad generalization of the models, which was confirmed when comparing RMSE distributions, where the test median values were almost 3 times higher than the train ones (**Supplementary Figure 5H**).

To assess if this over-fitting problem has implications on model performance across *DS*, potentially introducing bias in our results, we test the models against synthetic data. This data was generated by using LHS method, ensuring that the sample is representative of the variability within *DS* (details in MM section 4.2). For each synthetic design of each bioprocess, we compute the squared error (SE) using *in silico* validation with iJN1462. To assess the overall performance of each model, the range-normalized square root of the mean SE (NRMSE) was computed (**Figure 4A**). This metric provides a standardized measure of the prediction error relative to the range of the observed data, allowing for comparisons across different datasets and models. In this case, the results clearly indicate that the models do not perform good in synthetic data, having a median NRMSE of 58.0% of the target range. Also, when the individual SEs are plotted against true values, models generated from gcFront data cannot properly distinguish non-viable designs from high-performance ones (**Supplementary Figure 6**). This translated into having designs with a predicted *CY* as high as 0.89 that when validated are non-viable designs. Moreover, the trend in the SE distribution suggests that higher errors are made for low true values, indicating a bias toward designs with good performance.

**Figure 4.**
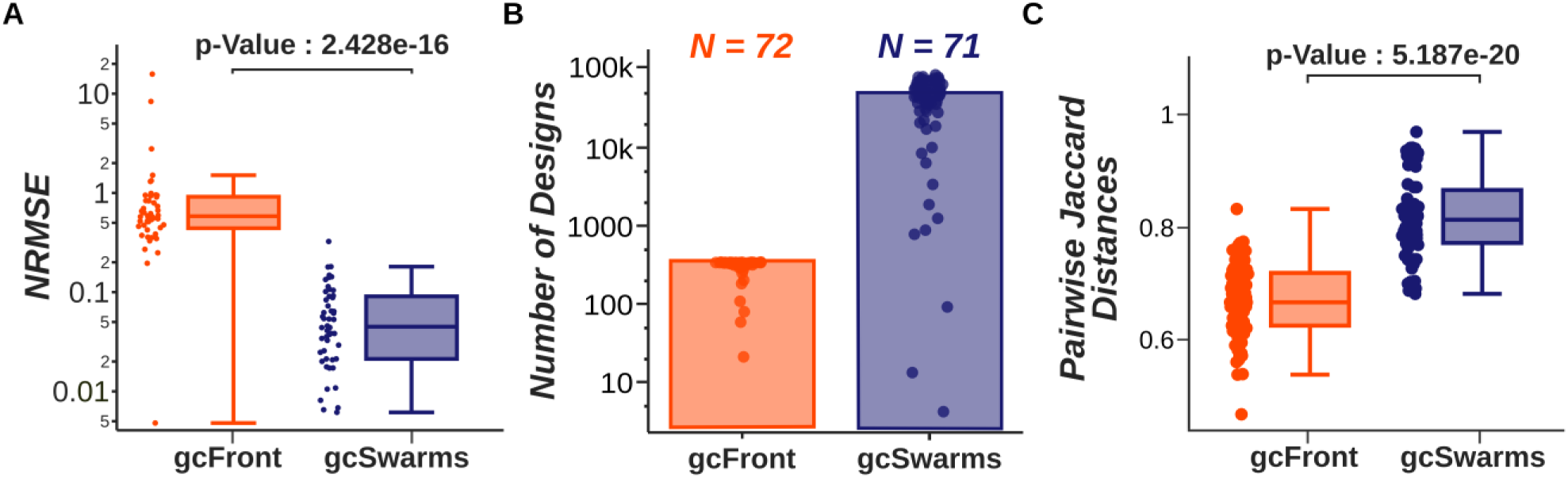
Comparison of gcFront and gcSwarms libraries for generating ensemble models. (**A**) Bar plot which recapitulates range-normalized RMSE values (NRMSE) of each of the ensemble model synthetic data predictions from gcFront (orange) and gcSwarms (purple). (**B**) Histogram showing the average number of viable SDs per gcFront and gcSwarms run. (**C**) Box plot of the computed median pairwise *Jaccard* distances of designs for gcFront and gcSwarms. For all plots, dots indicate NRMSE values of individualmodels, while the p-Value (when indicated) refers to the significance value obtained after a Mann-Whitney U test performed between data of different SD methods tested.

To analyze the causes of bad model performance, we inspect the generated SD libraries. The first thing that becomes clear is that gcFront only returns as a result libraries generated only by pareto optimal designs^10^, which explains the bias we observe in the model response. Furthermore, the library size is extremely small relative to the overall design space formed by approximately 100 candidate reactions (2) . This limited coverage makes it difficult to obtain a representative population of designs.

We next evaluated whether the design patterns within each library are sufficiently diverse and assessed the similarity between individual designs using two complementary metrics: a frequency-based measure (Shannon entropy per position) and a distance-based measure (median pairwise Jaccard distance). For details on the computation of these metrics, refer to MM 4.1. The results reveal substantial variability across bioprocess-specific models for both metrics, indicating that design diversity is highly dependent on the target bioprocess (**Supplementary Figure 7** and **Figure 4C** respectively). Overall, while some SD libraries exhibit high values for both diversity indicators, the majority are characterized by considerable homogeneity.

This is evident from the median values of computed metrics. Specifically, a median Shannon entropy per position of 0.532 suggests moderate diversity—positions in the binary design vectors are neither fully random nor strictly deterministic. In practical terms, this implies an imbalanced but not entirely skewed distribution of 0s and 1s. Also, Jaccard distance results indicate that designs within a library are relatively homogeneous (**Figure 4C**), having a median pairwise distance of 0.67. Notably, 38 out of the 72 generated libraries fall below this threshold, indicating frequent pattern recurrence among designs. This hampers the model generalization, as it is only learning from a small and not enough diverse subset of *DS*.

In summary, the lack of negative examples (designs with *CY* =0) added to the homogeneity of the design patterns that gcFront provides, result in an over-fitting of the ensemble models to the training data. Consequently, models perform poor quality predictions, making it necessary to find an alternative way to generate the initial library from an SD algorithm.

### 4 gcSwarms. A high-throughput strain design algorithm providing a ML-ready library for ML4SD

To deal with the problem of generating a design library having both design and performance data diversity, we propose a novel algorithm called gcSwarms. This algorithm consists of a Binary Particle Swarm Optimization (BPSO) approach specifically designed for metabolic engineering design optimization based on deletions (**Figure 5**).

**Figure 5.**
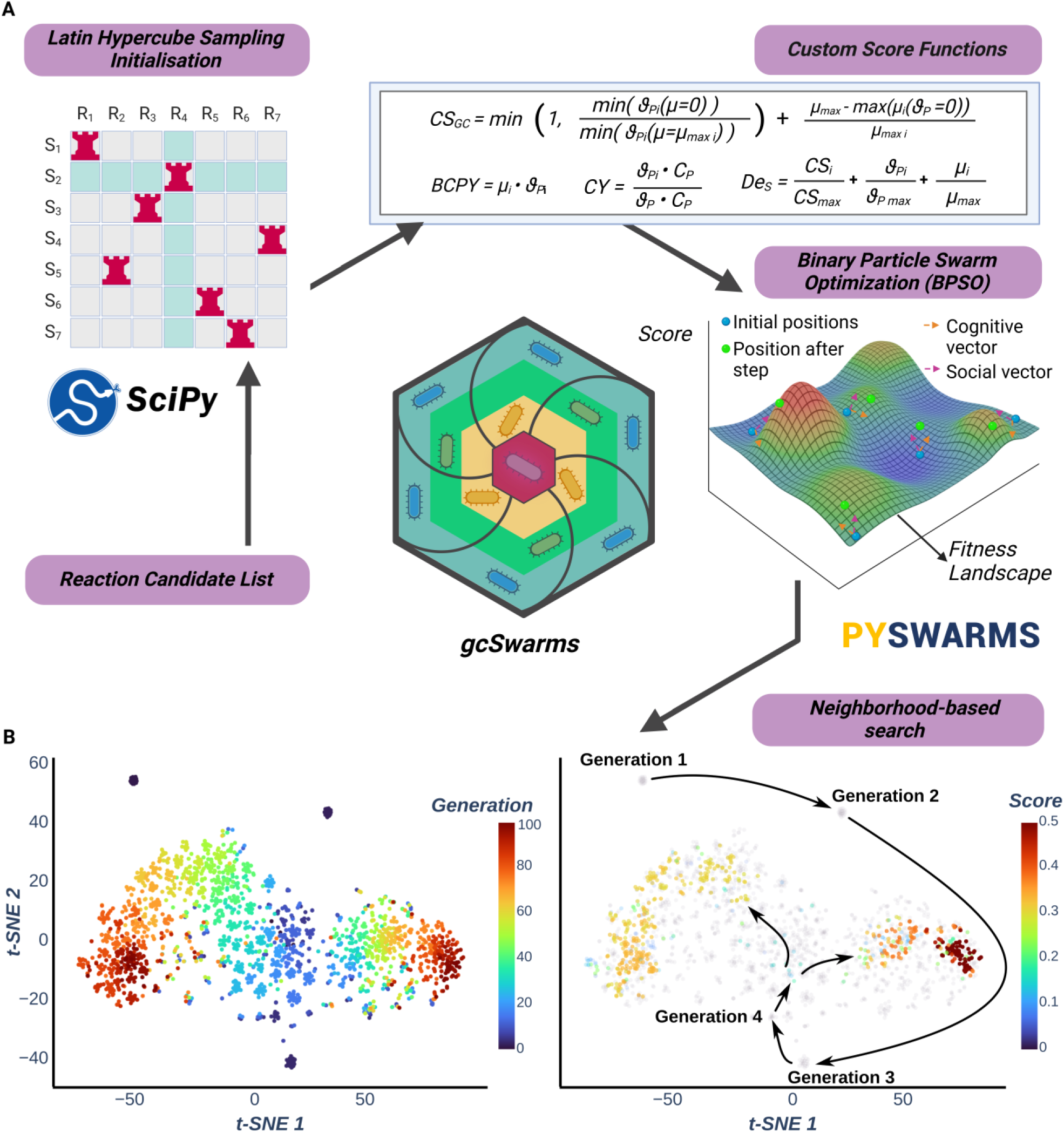
Overview of the gcSwarms search process for metabolic design optimization. (**A**) Main features of gcSwarms. Each particle in gcSwarms represents a strain design encoded as a binary vector of the same length as the candidate list provided. Initial population of designs is initialized using Latin Hypercube Sampling (LHS) to ensure enough variability coverage of the search space. Fitness evaluation of those designs is performed via a GEM-based function, that can be user-defined or selected among the predefined functions available. gcSwarms iterates over 100 runs of the *pyswarms* implementation of a BPSO, being the swarm size dependent on the number of designs to explore. Between iterations, the design space is surfed through a neighborhood-based search to balance exploration and exploitation. All described features mitigate local optima traps while generating a diverse design population in a single execution. (**B**) Representation of a design search example performed by gcSwarms, exploring a total of 2000 designs using a neighborhood-based search with 2 different groups. Each dot in the graph represents a given design, embedded in the design space by using t-SNE over their binary vectors, with color representing generation (left) or fitness score (right).

PSO is a population-based evolutionary algorithm first proposed 2 decades ago^46^. It has become one of the most popular algorithms for solving multimodal optimization problems due to its simple implementation and effectiveness. Several variants have been applied in many real-world optimization problems, recently including metabolic engineering^47–49^. In PSO, each particle in the population is represented by its position in the problem multidimensional space, constituting a potential solution whose optimality depends on a given function. Each particle adjusts their trajectories and positions by learning from their own experience and the global experience of the population (i.e. swarm), helping each other to find a better search space and converge towards an optimum solution to the problem. However, the algorithm may suffer from the problem of premature convergence, meaning that it gets trapped into the local optimum easily especially when solving complicated problems. Considering this problem, a lot of work has been done, resulting in the development of different variants of PSO in the past decades. Those variants mainly focus on balancing the exploration behavior of global search with the exploitation behavior of local search^50^.

In gcSwarms, each particle represents a binary vector corresponding to a design and following (*Def*. 1). Designs could have up to *d* maximum deletions, set by the user. gcSwarms initialize the positions of those particles following a Latin Hypercube Sampling (LHS) method which ensures diverse particle distribution across the *t-dimensional* search space (MM 2.5.2). After this, gcSwarms is set up to run for 100 iterations, with a swarm size that depends on the number of designs that the user wants to explore (*N*). The search during those iterations is made following a neighborhood-based approach. In this way, the swarm is divided into groups (g), limiting each particle’s awareness to nearby neighbors, and thus hindering premature convergence by balancing the exploration and exploitation behavior of the algorithm (details in MM 2.5.1).

The optimality/fitness measure for each particle position in gcSwarms is given by a GEM-based function that characterizes the performance of a given deletion design. Importantly, we have provided gcSwarms with the ability to incorporate custom objective functions. The “*DesignScores.py*” file allows users to create GEM-based scoring functions, optimizing for specific criteria. Nowadays, there are 4 different objective functions, all based on GEM-based metrics of a given deletion design (see MM 2.5.3).

Finally, PSO-based algorithms are programmed to retrieve only the best solution they found for a given optimization problem. However, in our case all the intermediate designs are crucial to capturing deletion influence over the final performance metric. Therefore, we use as the output of gcSwarms the full search history recording. All the mentioned features of gcSwarms contribute towards efficiently exploring metabolic designs while mitigating the risk of falling to a local optimum, with the final aim of generating a diversified design population within a single gcSwarms run.

To generate the SD libraries from gcSwarms data (D*_Sw_*), we apply our in-house SD pipeline with the same setup described before over the same bioprocesses as executed in gcFront, only changing the SD algorithm. For the specific gcSwarms parameters used for generating D*_Sw_* please refer to section 2.5.1 of MM. As a result of this, we identified deletion strategies for 64 bioprocesses (51.2%), indicating that some of them were more challenging to address within our imposed computation time (around 40 min). Those libraries contain an average of 5.9·10^4^ unique designs meeting the performance criteria (**Figure 4B**). This is considerably more than the libraries generated through gcFront and therefore, to our knowledge, for any other SD algorithm. Notably, those designs were obtained within less execution time (**Supplementary Figure 8**). This is extremely important for a DBTL, as it enables a quick execution of the design step in the first cycle round. Importantly, the designs within libraries are also quite diverse, having a median Jaccard distance of 0.44, significantly higher than libraries obtained from gcFront (**Figure 4C**).

For each of the libraries, we generate a bioprocess-specific ensemble model using the same cross-validation methodology as before, only varying the ensemble search time (see section 3.3 MM). Once the training was done, we measured the predictive power of the generated models within the train and test sets models through their R^2^. Considering the first, the models show a median R^2^ of 0.936, indicating good predictive power (**Supplementary Figure 5E**). Also, the R^2^ values for the test set are quite good (median of 0.872). Importantly, in contrast with gcFront, R^2^ values do not vary significantly between the datasets (**Supplementary Figure 5F**). These results indicate a better generalization of the models, which was confirmed when comparing RMSE distributions, where the test median values were approximately like the train ones (**Supplementary Figure 5G, H**). For comparing the models with the ones generated through gcFront, we compute the predictive performance of them using synthetic data as previously described. The obtained NRMSE values were significantly lower than those obtained using gcFront. Specifically, the median value was 4.4% of the target range, indicating good predictive performance of the models (**Figure 4A**).

Overall, we have shown that gcSwarms algorithm can generate design libraries in the order of thousand designs within a relatively short interval of time. Thus, considering also the size of the libraries, the diversity of input data and the performance of the ensemble models, we conclude that this SD algorithm meets our criteria for being part of our active learning platform ML4SD.

### 5 Deployment of ML4SD for a DBTL-based strain design optimization concerning the upcycling of lignin-derivative 4-hydroxybenzoate towards Nylon-6

Once we have all the needed modules that construct ML4SD, we apply the framework on a case study to prove its benefits. For assuring that we are going to obtain designs for a bioprocess optimization we are going to select between the ones within the feasible growth-coupled production space in *KT*. Among those, the bioprocesses derived from 4-hydroxybenzoate (4-HB) pose a clear biotechnological interest. This is because 4-HB is a major intermediate in lignin degradation pathways. Lignin is a complex and recalcitrant aromatic polymer found in plant biomass (**Figure 6A**). Its structural heterogeneity and resistance to degradation have traditionally limited its valorization in biotechnological applications. As lignocellulosic material constitutes on of the major industry subproduct, the efficient bioconversion of lignin-derived aromatic compounds into industrially relevant chemicals is a key objective in the development of sustainable biorefineries^51,52^.

**Figure 6.**
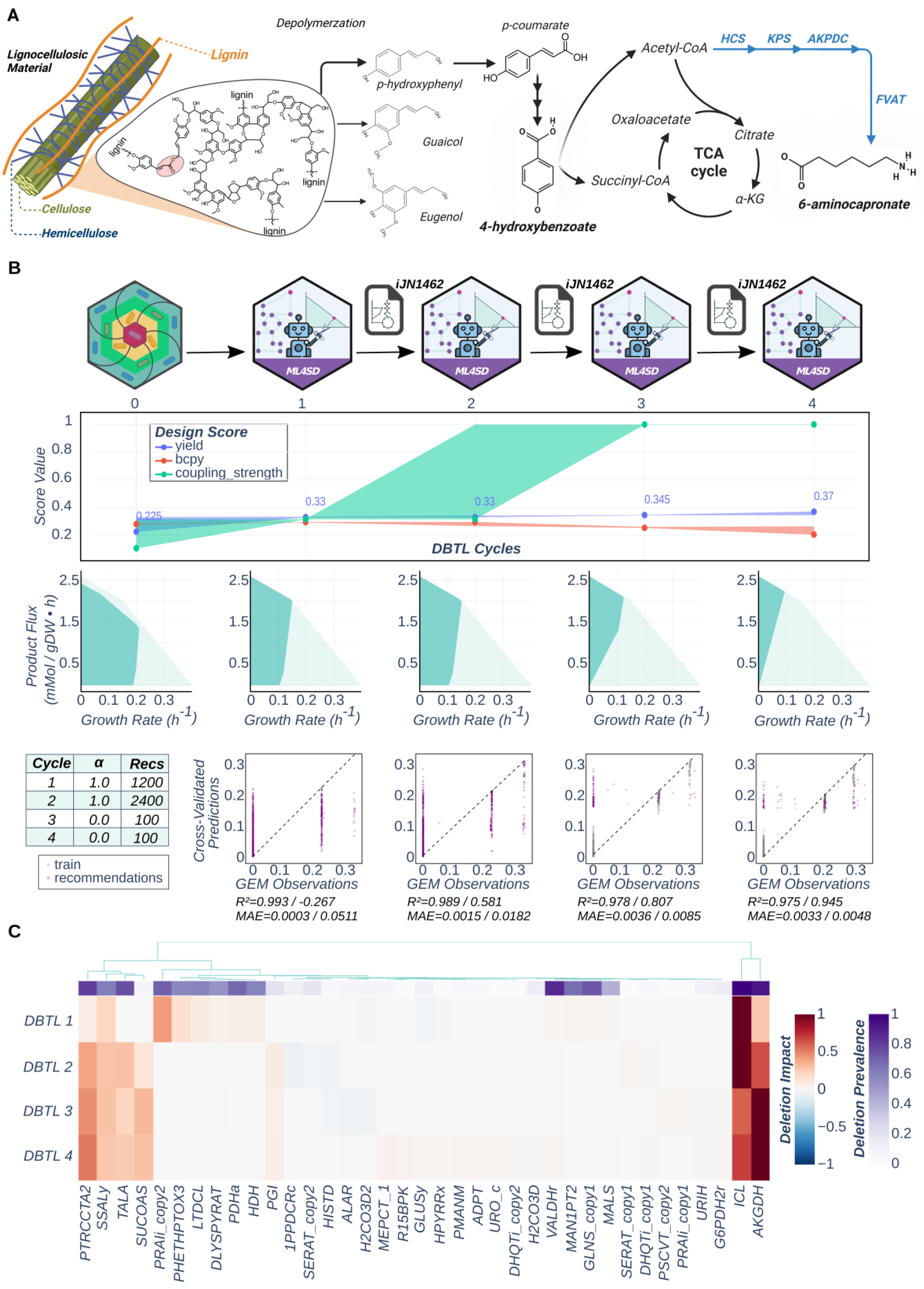
Results of an example of a lignin valorization DBTL campaign optimizing design carbon yield. (**A**) Scheme representing the natural source of the selected carbon source and its transformation into the nylon-6 precursor 6-aminocapronate. Reactions not included in iJN1462 are colored in blue. (**B**) Analysis of ML4SD results for strain design CY optimization campaign. Progression of fitness scores of best designs across DBTL rounds for 3 different replicates are shown (top). The area represents the range of fitness scores between replicates, while the dots represent the replicate 2 of CY campaign. Below, the envelopes of the best design (dark blue) for each round are represented in comparison with the WT (light blue). At the bottom, a summary table recapitulates the DBTL parameter setup chosen for the present case study. Following this, predicted cross-validations are plotted against true values according to GEM simulations for each DBTL round. Below each graph R^2^ and MAE values are shown. Those correspond to ensemble model predictions over the library used as training dataset (left) and over predictions of the library after all round recommendations have been added (right). (**C**) Clustered heatmap recapitulating importance and prevalence of deletions according to ensemble models of each DBTL round for the second replicate of CY campaign. The first parameter is computed from Shapley values of the top percentile designs within the generated library, while the second is the percentage of those designs in which a deletion is present (see MM section 5.2).

As previously mentioned, *P. putida* possesses an extensive enzymatic toolkit that enables the breakdown of a wide array of compounds that are challenging for other bacteria, including lignin-derived aromatic compounds^34^. Through specialized catabolic pathways, such as the β-ketoadipate pathway, these compounds can be converted into central metabolic intermediates, facilitating their transformation into value-added bioproducts. Moreover, this bacterium exhibits high tolerance to lignin-derived inhibitors, further enhancing its industrial relevance^53^. Advances in metabolic engineering and synthetic biology have further optimized *P. putida* for enhanced bioconversion, enabling the production of bioplastics such as polyhydroxyalkanoates (PHAs) and other commercially relevant metabolites^38,54,34^.

Among all feasible bioprocesses from 4-HB, we select the biosynthesis of 6-caprolactam for our study case. This molecule is the monomer for nylon-6, a widely used engineering plastic with applications in textiles, automotive components, and packaging materials^55^. Thus, the optimization of this bioprocess potentially boosts the valorization of lignin waste streams into high-performance biomaterials. Also, the traditional synthesis of nylon involves petrochemical routes, which are associated with high energy consumption and environmental burdens. Therefore, its replacement for a microbial conversion has great interest in reducing our reliance on fossil-based resources, aligning with the principles of circular bioeconomy.

To guide the strain design for this task, we use the iJN1462 high-quality GEM of *KT* as the digital twin of its metabolism. As ML4SD merges active-learning algorithms with a DBTL cycle methodology, it should allow an effective search of the design space based on knowledge gained in previous experiments. Following this reasoning, the initial library can be relatively small as long as it contains designs with a sufficient range of score values to learn. Moreover, the small size of the initial library is also desirable to reduce time and costs during the test phase in a wet lab setup. In such a scenario, the initial design library is going to be limited to the available time and resources of the lab. However, as we are running an *in silico* proof of concept, we have decided to set up an initial library size of 1200 designs, which is relatively small in comparison with the ones generated previously during the evaluation of the SD algorithms but large enough for gcSwarms to find relevant growth-coupled designs in most of the cases.

Another important aspect to consider is the target score to learn. Above all, our interest resides in the possibility of experimentally testing *in silico* predictions made for a given bioprocess for the ensemble model, enabling it to correct the GEM biases. In that respect, among all the scores available in gcSwarms, we have chosen to perform one campaign for carbon yield and another for BCPY, as both parameters can be estimated in a wet lab facility.

For the generation of an ensemble model of enough quality during the DBTL rounds, we need to consider an appropriate initial training time. We do it by following the same methodology as in section 3.3 of Materials and Methods. Importantly, as the recommendations of one cycle are added to the library, the training dataset increases with the number of cycles. Thus, we also decided to dynamically update the training time each cycle, maintaining the same data to time ratio.

For balancing the explorative and exploitative modes for computing *S(*D) within ML4SD, we use the parameter *α* (3). We consider ML4SD to be in explorative mode while *α*>0.5 and exploitative in any other case. Noteworthy, α also determines the importance of each deletion feature during the computation of the ML-based sampling distribution, that we discuss in section 1.1 of MM. As there is no clear guidance for the selection of this value, we adopted a strategy of fully exploring, fully exploiting. This translates into selecting an *α* value of 1 for the two first cycles and a value of 0 for the last cycles (**Figure 6B bottom-left**). This yields two explorative cycles, focused on producing recommendations that explored regions of the design space where the predictive power of the model is poor, followed by two exploitative cycles where recommendations are made based on their predicted potential for score optimization.

Concerning the number of recommendations to make on each cycle, we took guidance from a study which compares the predictive power of a ML model across different DBTL cycle strategy scenarios^56^. Along all the tested scenarios, no significant differences for the same number of data points could be observed. However, an incremental strategy is the one reaching the maximum predictive power within the minimum number of cycles, so we decided to adopt it in our workflow. We slightly adjust the strategy, starting with a small initial dataset that increases proportionally to the number cycles while the DBTL mode is in exploration mode. In contrast, for the exploitative mode we decided to add a fixed number of recommendations because longer computing times are needed for generating them.

Regarding the performance score to optimize, we use those available in gcSwarms that are easily measured from a biorefinery facility, specifically *CY* and *BCPY* ^(^4^)^ and eq. ^(^5^)^. *CY* is useful when maximum conversion of prime material is required. This can be the case either when using a costly substrate or for bioprocesses with tight sustainability goals. The second applies for our target bioprocess, as its implementation would reduce CO2 emissions bypassing petrochemical production of nylon, being therefore crucial to optimizing carbon utilization. On the other hand, *BCPY* is also a relevant measure as it accounts for the chassis growth in addition to the target production. As we are looking for designs yielding production coupled to growth, when *BCPY* favors designs leading to higher biomass formation, it also is increasing target production, ending with a more balanced search of designs.

The results obtained in both campaigns show that across different DBTL rounds, designs leading to higher scores are found. In the case of the *CY* campaign, the higher increments on this parameter used to happen in the first rounds of the DBTL. This increase, although less pronounced, continues throughout all DBTL rounds, reaching a *CY* value up to 164.4% respect the best design in initial library. Interestingly, the increment of *CY* also implies an increment of *CS* in central rounds of the DBTL. This change to strong coupling can be noticed in the envelopes resulting from the best recommended designs for the 2^nd^ or 3^rd^ round. Moreover, this phenotype is always found in all the recommendations made in the exploitative rounds of DBTL across the different replicates, highlighting the relevance of this algorithm mode for finding high-performance designs (**Figure 6B middle**). Concerning the active-learning process, the model learning only from gcSwarms library has a very low predictive power over the final library, while the model in later rounds progressively improve it (**Figure 6B bottom**). This implies that new knowledge is incorporated through the DBTL rounds that enable the models to generate better recommendations. This applies in each replicate of the campaign, remarking the reproducibility of the active-learning methodology (**Supplementary Figure 9,10**). This is also supported by the importance that ML models give to different reactions across the DBTL rounds.

For all replicates, the initial ML model shares some of the most important deletions with the rest of the models but differentiates for a few targets that can be either learnt or forgotten in following rounds (**Supplementary Figure 11**). An example of both processes can be observed for the 2^nd^ replicate. In this case, the deletion of PTRCTA2 has no importance at the beginning but it gradually gains importance through the rounds, becoming the 3^rd^ most important deletion at the end of the DBTL. Conversely, the deletion of PRAli_copy2 reaction is given considerable importance for the first-round model but not for the rest (**Figure 6C**). Also, analyzing those relative importances of deletions is useful for prioritizing targets for the *in vivo* implementation of designs. Importantly, we found a minimal intervention strategy derived from the 3 most important deletions across all replicates for this campaign (ICL, AKGDH and PTRCTA2/SSALy). This deletion motif recapitulates mechanistical information of the GEM, as it is the minimal modification to redirect the flux from the TCA towards 6-caprolactam via increasing flux through acetyl-CoA (**Supplementary Figure 12**). We consider that all these findings prove the potential of the active-learning methodology for guiding strain design tasks.

Results for the *BCPY* campaign also remark the framework ability to improve initial designs provided by gcSwarms, leading to designs with better performance in all the replicates. Also, ML models within each replicate follow the same pattern in the predictive power as described for *CY* campaign, proving that the active-learning methodology incorporate knowledge through the DBTL rounds properly regardless of the parameter used (**Supplementary Figure 13-15**). This is corroborated through the analysis of the importance each model gives to deletions (**Supplementary Figure 16**). Interestingly, the 3 deletions that form the minimal design are also within the most important deletions across all replicates in *BCPY*. This suggests that this design is a shared motif across campaigns leading to more specialized designs improving each of the parameters used.

Despite the positive results obtained from both campaigns, there are significant differences between them. Notably, the maximum increase in design performance in BCPY represents only 101.6% of the best design found in the initial library. However, this can be explained by the fact that for the BCPY case, gcSwarms is able to generate in the initial library a design with a BCPY of 92.7% of the theoretical maximum for this bioprocess. This hinders the potential of ML4SD when compared to the *CY* campaign, that has a greater range of exploitability as gcSwarms reaches between 52.5% to 77% of the theoretical maximum (**Supplementary Figure 17**). Also, if we compare both campaigns in terms of the generation of high-performance designs, 53.6% of viable designs obtained across the 3 replicates of *CY* campaign are within the exploitative range, in contrast with 15.1% on the *BCPY* one (**Figure 7A**). Therefore, we can conclude that ML4SD conduct a data-efficient search of strain designs, which is more advantageous when SD algorithms provide designs with a performance far enough from the global maximum.

**Figure 7.**
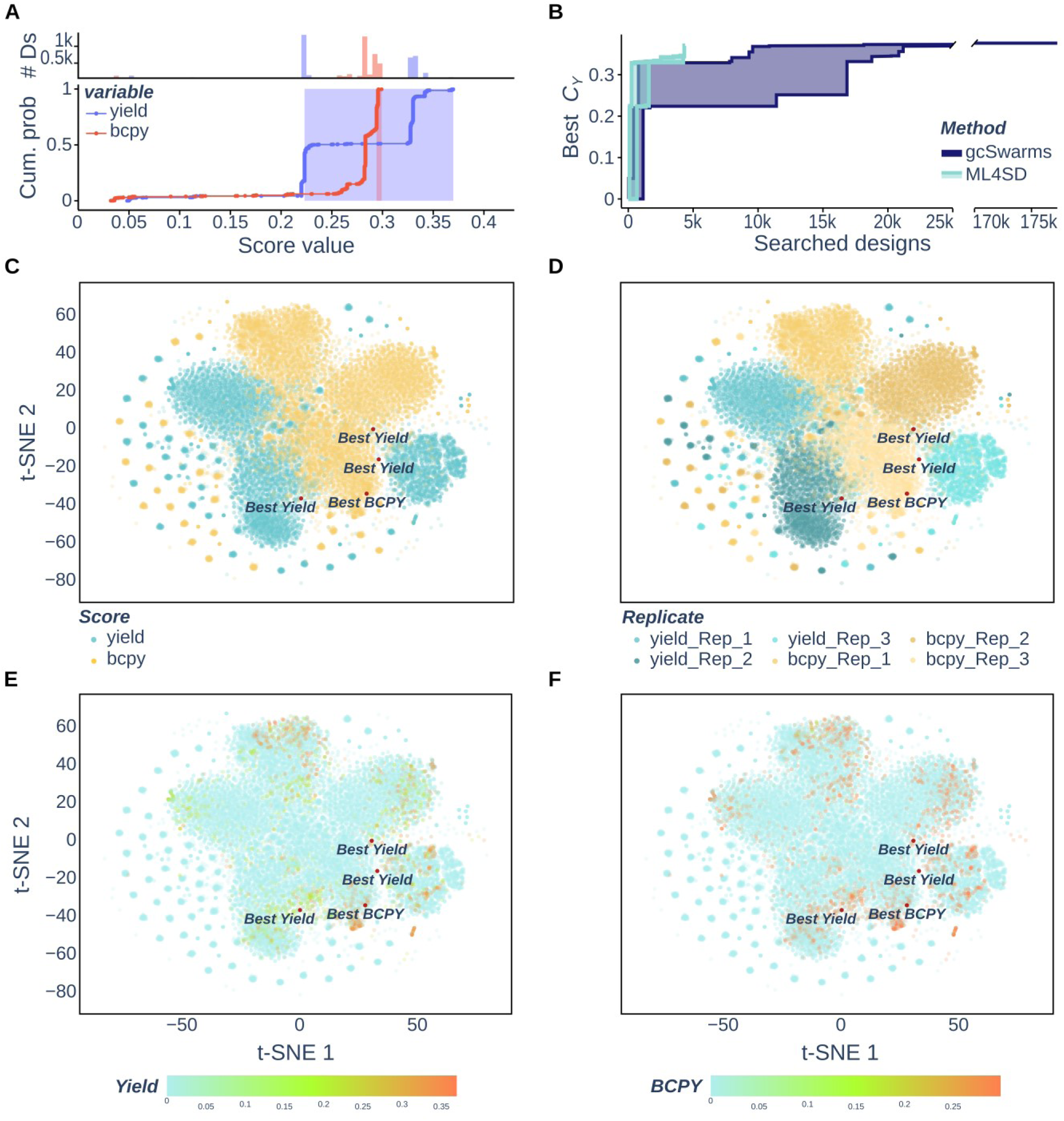
Design Space exploration through BCPY and CY campaigns. (**A**) Cumulative distribution plot of viable designs found among 3 ML4SD replicates for each campaign. The colored area represents the exploitative range, which is the difference between the best design score of *gcSwarms* and the best design found in the final library. Above, a histogram counts the number of designs (*# Ds*) across different score values. (**B**) Data efficiency plot comparing ML4SD and gcSwarms. Both areas represent the range of the best *CY* values found given the number of explored designs across 3 replicates of ML4SD (light blue) and gcSwarms (purple). (**C-F**) Representation of the design space explored across all campaign replicates. Each dot in the graphs represents a given design, embedded in the design space by using t-SNE over their binary vectors, with color representing campaign (**C**), replicate (**D**) or the target scores to optimize (**E, F**).

Data efficiency is key when considering using MLS4SD instead of simply a SD algorithm. Thus, to check the advantage that our active-learning methodology has, we track the best performance along with the number of explored designs for 3 gcSwarms libraries generated with the setup used in the comparation with gcFront and the 3 replicates of *CY* campaign (**Figure 7B**). We found that ML4SD reaches its maximum value by searching through 4316 designs. In contrast, gcSwarms took between 10869 and 30615 designs to reach a similar *CY* value, which is from 2.5 to 7.1 times more data depending on the replicate. Importantly, for generating the gcSwarms library we used 32 cores within a HPC server for 44 minutes, which is considerably resource intense. On the other side, ML4SD was run within a laptop setup (***Table 4***) using 6 cores for the initial library generation, taking an average time of 74 minutes to generate the final library containing the designs of all DBTL rounds. Consequently, ML4SD proves to be more resource-efficient in comparison with gcSwarms, with the capability of generating high-performance designs within a few minutes using standard hardware equipment.

Finally, to analyze the exploration carried out by ML4SD, we structurally characterize all the generated libraries. For this, we compute the design space by performing the same dimension reduction methodology followed with the metabolite fingerprints (see MM section 1). The resulting design space shows that designs from the different campaigns share a commonly explored region, supporting our hypothesis that final designs derive from simpler shared motifs (**Figure 7**). Despite this, all replicates in each campaign explore unique zones of the design space, indicating that it is convenient to run more than one DBTL cycle to maximize its coverage (**Figure 7D**). Interestingly, replicates of the same campaign tend to be closer in the design space, suggesting closeness of design patterns within a campaign, which is indicative of a specialized search depending on the parameter to optimize. However, as the 2 parameters explored here are closely related, both campaigns end to explore design space regions where there are also optimal designs for the other parameters. Moreover, if we look closely to the design space with the mapped values of *BCPY* and *CY*, we can see that all the best designs are in regions between both campaigns and are relatively close between them, pointing to a convergence on the search of pattern designs across campaigns (**Figure 7E, F**). Counterintuitively, those designs are not necessarily on regions with high density of high-performers, indicating that the design space consists of a rugged landscape typical of a complex genotype-phenotype mapping. This highlights the importance of tools like ML4SD, that will aid the metabolic engineering community to surf over this space to generate useful strain designs for complex metabolic engineering tasks within minimal time and computation resources.

Overall, we can conclude that the application of ML4SD workflow over the upcycling of 4-HB resulted in recommendations optimizing initial design libraries for BCPY and *CY* campaigns. From those designs a minimal pattern could be retrieved from the ensemble models by using SHAP values, which remarks the utility of such models not only for generating recommendations but also for recapitulating design principles. Importantly, those recommendations were obtained in a data-efficient way, significantly improving the results from SD algorithms, which is crucial for metabolic engineering tasks. Altogether, those results highlight the potential of ML4SD to contribute to the development of biofactories by efficiently guiding their search toward optimal strain designs.

## Discussion

In this work, we developed a novel, diverse-search knockout algorithm, gcSwarms. We embedded it within an active-learning DBTL cycle, ML4SD. This active learning methodology was thought to guide SD campaigns through the optimization of deletion designs using custom GEM-based scoring parameters. We applied ML4SD to improve growth-coupled designs for converting lignin-derived 4-hydroxybenzoate to 6-caprolactam in Pseudomonas putida.

For this novel approach, we initially selected gcFront for library generation. However, gcFront returned only Pareto-optimal knockouts^10^ and therefore small, homogeneous libraries that lack non-viable examples, on which ensembles overfit. gcSwarms, with its LHS initialization and keeping the full search history, favored larger and more diverse libraries, enhancing the same ensembles learning. Under a similar pipeline and time budget, gcSwarms also returns more unique designs in less time than gcFront, critical for having a more representative set of the design space.

However, during the library-generation screen utilized to assess the ensemble performance across a range of bioprocesses, neither gcFront nor gcSwarms generated a library for each bioprocess within the specified search criteria. While this can simply mean that some bioprocesses couplings are not possible, the fact that different bioprocesses were determined feasible by one of the algorithms suggests that more time should be given to the SD algorithm to correctly generate the initial SD library without false negatives.

Regarding ML4SD, other active-learning DBTL has already raised titers, but those approaches consist on recommending promoter combinations or multiplexed CRISPRi constructs from sparse in vivo data, testing recommendations *in vivo* within each cycle.^14,57^ Gene deletion is experimentally demanding and therefore another type of strategy should be followed. DeepGDel^26^ infers binary growth-coupled strategies from curated sequence databases, which limits its scope, and make one shot predictions without metabolic topology or a production metric. Therefore, we decided that ML4SD should sits between these lines: ensembles are trained at GEM-scored knockout libraries and recommend the next deletions. However, is through SHAP values that we recover design rules rather than only a next improved design. Consequently, we recommend running the first ML4SD cycles only in silico, scoring recommendations on the complete reconstruction. Once the last ensemble is explained, in vivo test strains should be constructed from the highest-importance deletions rather than from every recommended vector. That sequence defers wet-lab effort until the motif is stable, instead of reconstructing a library every round.

Considering that, we successfully applied ML4SD on a biotechnologically relevant bioprocess: the transformation of 4-hydroxybenzoate to 6-caprolactam, a nylon precursor. We improved initial maximum yield up to 164% for one campaign, identifying a potential minimal design for growth-coupled production of 6-caprolactam. Additionally, we observed an interesting pattern from the other campaign: when an initial swarm search already approaches the theoretical bound, as for biomass-coupled yield, there is little headroom for the learner. This raises questions concerning DBTL setup parameters, including the optimal scoring variable to select, the initial library size or the number of recommendations to make, whose answer will require extensive proofs.

This results remain a computational proof of concept and highlight the reason why a strain-design DBTL that is capable of ingesting both in silico libraries and later in vivo scores is of great importance: GEMs provide stoichiometric bounds and a warm start when there are no existing titers; measured production then corrects kinetics, regulation, and genetic-tool failure that flux balance cannot see. Without that second stream, recommendations remain ranked lists on a digital twin rather than designs that have been shown to yield a product in the host.

Inevitably, this leads to the immediate next step being the construction of high-importance knockouts, especially the minimal design identified in this study. Feedback from those experimentally scored strains (either by carbon yield or BCPY) should then be incorporated by transfer learning, using the outputs of the individual models in the final ensemble to fit the experimental data. Finally, a further extension would be made to include additional reaction and metabolite information to the training data. Descriptors of substrate and product structure, along with topological reaction data would let the models learn how growth-coupled knockout patterns relate to metabolic context, improving deletion deign prediction.^23,26^

Converting lignin-derived aromatics into nylon monomers is a circular-economy target that still depends on competitive titers, rates, and yields. By pairing a diverse in silico design search with an interpretable active-learning cycle, ML4SD offers a practical route to propose—and later test—growth-coupled strains for that class of bioprocess needed to redesign our economies.

## Materials and Methods

### 1 Representation of chemical diversity within iJN1462

To address the chemical similarity of the metabolites in iJN1462, we firstly use the *PubChemPy* python package ( https://github.com/mcs07/PubChemPy ). With it, all metabolites were mapped against PubChem database^58^, extracting the isomeric smiles for all the matches (for details see *get_metabolite_smiles.py*). With those, all fingerprints were computed using the *GetRDKitFPGenerator* default class of the *RDKit* python package version 2022.03.5 ( https://doi.org/10.5281/zenodo.6961488 ).

Fingerprints are used for generating the chemical space of iJN1462. This is done by embedding them in a bidimensional space through the t-SNE implementation in *Scikit-learn*^59^ with PCA initialization for preliminary dimension reduction. To select a suitable embedding, we evaluate each of them through specific performance metrics along with clustering metrics of the restructured data.

Specifically, for each tested perplexity value of t-SNE, DBSCAN was run over the generated chemical space through a range of eps (from 5 to 50) and minimum fingerprints per cluster (from 2 to 10). Through all possible parameter setup combinations, we select the one yielding the higher Calinski–Harabasz index (CHI) for assessing the data modularization. Thereafter, for each generated chemical space, we compute use the average fraction of 10^th^ nearest neighbors in the original high-dimensional data that are preserved as 10^th^ nearest neighbors in the embedding (KNN ratio) to quantify the preservation of the microscopic data structure^35^. Nearest neighbors were computed by using Tanimoto pairwise distances in the case of fingerprints, while Euclidean pairwise distances were used in the case of embedded fingerprints.

The final embedding was selected following this criterion:

1) The embedding-specific clustering must not have outliers
2) The KNN ratio must be above 0.5 for assuring well represented local similarities
3) Among parameter setups meeting previous conditions, select the one with highest CHI

The evaluation metrics of all clustering and embeddings can be seen in **Supplementary Figure 1**. The selected embedding was the one resulting from a t-SNE with a *perplexity* of 45 and a DBSCAN with a *eps* of 5, an a minimum number of fingerprints per cluster of 2.

For the iJN1462 code implementation of metabolite smiles retrieval and the generation of the chemical space, please refer to files *“get_metabolite_smiles.py”* and *“GEM_chemical_space_representation.ipynb”* respectively, both within the code folder.

### 2 Strain-Design methods

#### 2.1 Generation of configured GEM

In the present work, we use *iJN1462* as the base GEM. However, this model does not comprise all reactions needed for the generation of all tested products, which is a common problem during strain design. To overcome this, we enable the use of a repository model harbouring all needed reactions. In this work we use a GEM derived from *IJN1411* generated in a previous study^39^. Then a reduction of the model is carried out for each bioprocess to speed up the search of deletion designs through strain design algorithms. As it can be seen in **Supplementary Figure 3**, the reduction consists of the elimination of biomass reactions other than the one that has been assigned as a target, along with reactions that do not carry any flux (blocked reactions). Reactions considered to be blocked are those not carrying flux when doing a loop-less FVA with a minimum growth of 20% in comparison with WT. All those steps were done over the GEM growing in minimal media using only the carbon source indicated by bioprocess specifications, with an uptake rate (*US*) fixed to a carbon flux of 36 mMol/h. This model is henceforth used to generate both the strain designs and the candidate reaction list.

#### 2.2 Selection of candidate reactions

To reduce the computation times of both gcFront and gcSwarms, we used a list of candidate reactions obtained from the GEM (RC). For constructing such a list, firstly we exclude from the GEM all essential, transport and no GPR associated reactions. With the obtained candidates, a thorough filtering was made by imposing a carbon limit filter to exclude from the list those reactions whose main reactant has a number of carbons higher than 12. The rationale behind this step is that those reactions are considered to be out-of-the-core (peripheric) reactions that have no significant impact on GC strategies, with little flux passing through them^9^. Ultimately, a reaction lumping was made by grouping reactions that are constrained to carry equal flux due to metabolite stoichiometries, selecting one reaction as the knockout representative of the group. With all those steps we achieve a RC on the order of one hundred candidates (varying with the specific bioprocess), therefore reducing the design space considerably (from around a thousand reactions of the reduced model) and optimizing the SD search.

#### 2.3 MCS analysis for coupling test

This step was executed before the execution of the SD pipeline as a filtering step of bioprocesses that are not feasible under the selected design criteria. The analysis was based on a modification of the pipeline performed in a previous work using minimal cut sets^41^. This analysis was implemented with the StrainDesign python package^60^. The entire workflow can be found in the script “*Strain-Design-Pipeline_feasibility_putida.py”* within the code folder.

For each coupling bioprocess candidate, the following steps are performed:

1) The original GEM is processed as described in Section 2.1 of materials and methods to generate the reduced model.
2) The product exchange reaction is added if not already in the reduced model. Such reaction is maximized in the GEM via solving an appropriate linear optimization (LP) problem. With this value, the maximum molar product from substrate yield 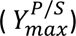 is computed and the minimum yield required for the bioprocess 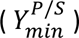 is set as one percent of 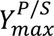. If the maximum flux of the product is zero, then the metabolite cannot be produced at all, and the forthcoming steps are not applied.
3) The growth prediction for the reduced model is made via solving an FBA with the core biomass reaction of the GEM as the optimizing objective. From this value, a minimum biomass from substrate yield 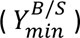 is set to 0.01 gDW mmol substrate^-1^.
4) A maximum knockout limit (*d*) of 30 deletions is set.
5) The candidate reaction list is computed as described in section 2.2 of materials and methods.
6) The constraints for the MILP are constructed by imposing both 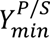 and 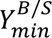 as the minimum product flux and growth respectively.
7) The MILP is run with a time limit of 10 min. As said above, the selected implementation of the MCS was the one implemented by the *StrainDesign* python package with *Gurobi* as the selected solver.

Specifically, the function *compute_strain_designs* from the mentioned package is run using the parameter setup in ***Table 1***.

**Table 1.**
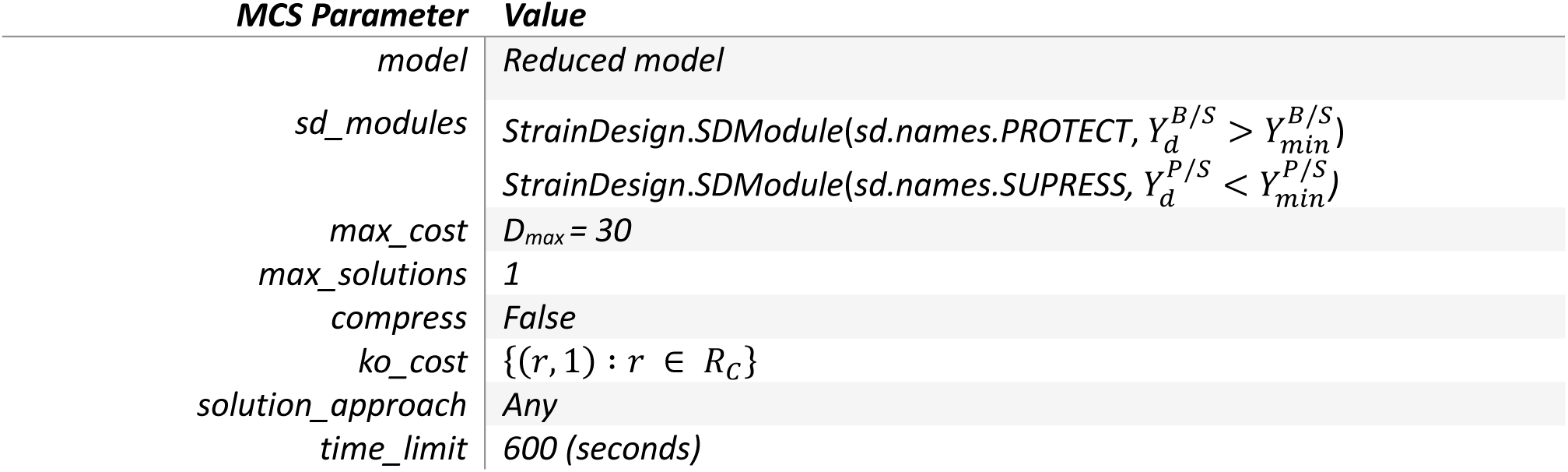
Parameter setup for running the MCS test.

| **MCS Parameter** | **Value** |
| --- | --- |
| *model* | *Reduced model* |
| *sd_modules* | *StrainDesign.SDModule(sd.names.PROTECT, $Y_d^{B/S} > Y_{min}^{B/S}$)*<br>*StrainDesign.SDModule(sd.names.SUPRESS, $Y_d^{P/S} < Y_{min}^{P/S}$)* |
| *max_cost* | $D_{max} = 30$ |
| *max_solutions* | 1 |
| *compress* | False |
| *ko_cost* | $\{(r, 1) : r \in R_C\}$ |
| *solution_approach* | Any |
| *time_limit* | 600 (seconds) |

Using this setup, we will search for MCS until a solution is found, the time limit is reached or when the problem is determined by the solver to be infeasible (meaning that coupling is not possible).

Those steps are applied up to 10 times (**C**) for each selected bioprocess, using a different solver seed for each execution, which leads to a different exploration of the search space^41^. Finally, those bioprocesses who do not pass at least one test are excluded from the gcFront or gcSwarms execution to save computation time.

#### 2.4 gcFront setup

gcFront was run in Elbrus HPC setup (***Table 4***) using 32 threads for parallel computation. The software was executed in MATLAB version: 9.13.0 (R2022b).

For each bioprocess having at least one MCS passed test, the candidate lists computed as indicated in section 2.2 (**D**) were passed to gcFront to reduce its design space. Also, *d* was fixed to 30, considering this value as a maximum for experimentally feasibility within a reasonable time range. The runtime was set to 6 hours, as used for benchmarking in the original paper^10^. All non-default parameter values used for gcFront design search are compiled in ***Table 2***.

**Table 2.**
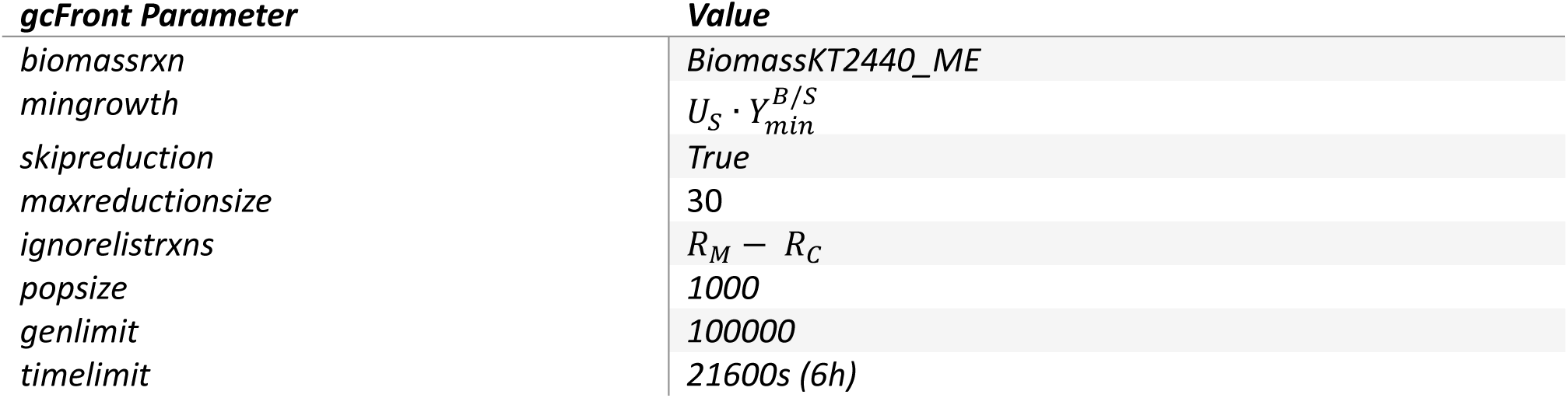
Setup of parameter values in gcFront that are different from default.

#### 2.5 gcSwarms algorithm

The gcSwarms algorithm was developed using Python language version 3.9.7 as the selected framework. Its implementation can be found in the files *“DesignScores.py”* and *“PSO4SD.py”* of the utils folder within ML4SD GitHub repository. Regarding the comparison with gcFront, it was run in Elbrus HPC setup (***Table 4***) using 32 threads for parallel computation. However, for the ML4SD study case was run in a laptop setup (***Table 4***) using 6 threads for parallel computation.

##### 2.5.1 Implementation of the Binary Particle Swarm Optimization algorithm

We use the BinaryPSO class of the *PySwarms* python package^61^ as the core of gcSwarms algorithm. It is important to note that 2 files of the source code belonging to *PySwarms* have been modified (*“base_discrete.py”* and *“binary.py”*) to enable the entire search history recording. This enables us to use all the intermediate designs found during the search as the result of the algorithm, instead of using only the best scored design as in the original implementation. To ensure reproducibility of gcSwarms algorithm, both modified files are included in the GitHub repository.

In gcSwarms, each particle position during the search is equal to a design defined as a binary vector. This vector has a length equal to the number of different candidate reactions (*t*) and with a maximum number of ones (*d*), both set by the user. Our implementation has a fixed number of 100 iterations, but the user can indicate the number of designs to search over (*N*), increasing then the number of swarms per iteration. In our case, we have selected a search over 180000 and 1200 particles for the gcFront comparison scenario and the

DBTL respectively. It is important to note that despite the user limits the search space by imposing *d* maximum deletions, the search space is still huge, with a number of combinations given by (2). As the gcSwarms search takes place in such space it is crucial to both initialize the algorithm with a diverse enough particle population and avoid falling in local optima.

To address such a problem, we have followed different strategies. Firstly, we initialize the PSO algorithm with a particle population generated by a design of experiment (DoE) methodology called Latin hypercube sampling (LHS; explained on section 2.5.2). LHS outperforms other methods such as random sampling in generating a diverse population that captures the variability of the search space, making it a good fit for our needs^62,63^. Secondly, we have used a dynamic neighborhood-based searching strategy in which the whole swarm population is dynamically divided into several groups (*g*) given by the user. In this way, each particle is only aware of the 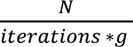 nearest particles to compute its cognitive and social vectors that will lead the particle to its new position, making it harder for particles to fall into a single local optimum (**B**). Such strategy has been used for approaching multimodal problems through PSO and has only recently been applied to optimize metabolite production using GEMs^49,50^. To facilitate balanced exploration and exploitation within each neighborhood, balanced values for cognitive and social coefficients are chosen (c1=2 and c2=2 respectively).

Finally, gcSwarms needs a function to guide the design search in a significant way. For this, a python class is implemented (see in *“DesignScores.py”*) that gives the user the capacity to create custom, GEM-based scoring functions of the designs. This enables the user to search for designs that optimize their task-specific criteria. However, we have implemented several scoring functions that are commonly used in the field (see section 2.5.3).

##### 2.5.2 Latin Hypercube Sampling for initializing search to promote design diversity exploration

The initialization of particle positions influences the search process in a neighborhood-based PSO algorithm by means of selecting a restrictive area within the whole search space^50^. To provide a representative enough initialization we applied the following steps:

1) Select the values for d and t for a given reaction candidate list of length t
2) Compute the particle population size per iteration (n = N/iterations)
3) Compute the number of particles for each possible deletion number as n/d
4) Apply LHS to sample n/d designs as lists of reactions for each possible deletion number from the candidate list
5) Transform all designs to binary space (see section 3.1)

##### 2.5.3 Implemented GEM-based objective functions

Although in the present work we only apply 2 different objective functions, in gcSwarms source code we implement a total of 3 commonly used objectives and 1 experimental score. All the objective functions described in this section are GEM-based. However, some of them (carbon yield and biomass-coupled product yield) can also be calculated experimentally *in* vitro. Those are useful for design validation and for learning purposes in ML. The implementation of all of performance metrics can be found in *“DesignScores.py”*.

**Carbon yield (*CY*).** For a given design *i,* this measure was computed following (4)

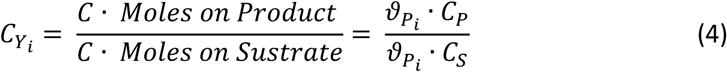

**Biomass-coupled product yield (*BCPY*).** For a given design *i,* this measure was computed following (5)

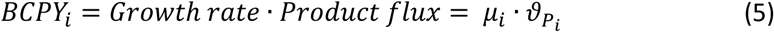

**Coupling strenght (*CS*).** This measure was implemented for the first time in the gcFront^10^. Following their implementation, for a given design *i,* we compute this measure according to (6), considering only designs whose implementation yields a product flux above zero. As the maximum value at both sides of the adding symbols can be 1, the maximum value of *CS* is 2, indicating the maximum coupling a design can have (*CSmax*).

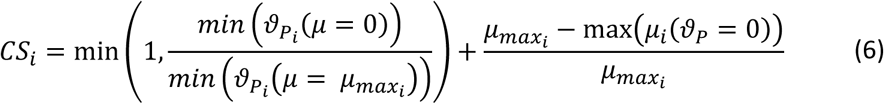

**Design Score (*DeS*).** This measure consists of the integration of other three that impact on the design suitability. Those are the growth rate (*µ*), the product flux (ϑP) and *CS,* informing about the biomass, the production and the degree of coupling between them respectively. It can be considered as a simplification of the multi-objective optimization made in gcFront^10^.

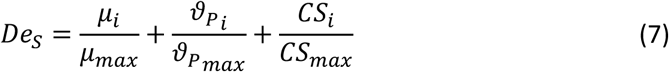

For clarification of the notation considering all equations described in this section please refer to **Supplementary Figure 4**.

### 3 Machine Learning methodology

The implementation of the different methodologies can be found in the file *“TrainSpecialistModels.py”* of the ML4SD GitHub repository.

#### 3.1 Library data preprocessing for model training

As previously said, in this study the strain designs are defined as binary vectors where 1 are reactions included in the design and 0 are those absent. For generating this data structure from a string of reaction deletions, we use the list of candidate reactions computed as indicated in section 2.2 of MM. With this bioprocess-specific list, the reaction deletions are one hot encoded into the final binary array that will be used as the training data for our ensemble model. It is important to note that this process is only applied to libraries generated by gcFront, as gcSwarms use this data structure as default output.

#### 3.2 Cross-validation methodology for ensemble generation

Although most of the ensemble model generation is automatic, some important decisions are left for the end user to make. This includes the cross-validation methodology, the training time left for the ensemble model construction and the individual models and the maximum model size among others. For our regressor implementation, we select the parameters in ***Table 3***.

**Table 3.** Selected values for cross-validation parameters of AutoSklearnRegressor class.

| Parameter | Value |  |
| --- | --- | --- |
| *time_left_for_this_task* | For gcFront: 600 s | For gcSwarms: 7200 s |
| *resampling_strategy* | "cv" |  |
| *resampling_strategy_arguments* | *train_size*: 0.8<br>*shuffle*: True<br>*folds*: 5 |  |
| *memory_limit* | 1024 Mb |  |

Concerning the validation method, with the selected set-up we have chosen a variant of k-fold cross-validation. In this method the dataset is divided into k folds. Each fold is used once as the validation set, while the remaining folds are used as the training set. This process is repeated k times (once per fold), and the model performance is averaged over the folds to evaluate its generalization capability. For further clarification, our specific steps of the validation process will consist of the following ones:

1) The entire dataset is shuffled.
2) The dataset is then split into 5 folds of approximately equal size.
3) During each iteration, a fold is used as the validation set, and the remaining 4 folds are combined to form the training set.
4) Of the 4 folds used for training, only 80% of those samples are used to train the model in that iteration. The remaining 20% of the training folds are ignored for that iteration.
5) The model is trained 5 times, once for each fold acting as the validation set.
6) The performance metric (R2 in this case) is computed for each fold, being the final performance score the average of the scores across all 5 folds.

This approach effectively acts as a form of nested resampling, balancing the need for both robust validation (via cross-validation) and smaller training subsets to ensure models generalize well and avoid overfitting^64^.

#### 3.3 Estimation of search time for ensemble generation

For the estimation of the optimal search time for ensemble, we have selected different values for the different sources of the design library data, as the volume and diversity of them are quite different. For estimating a reasonable time, we have assumed that the more data the more time needed for training and so, for each of the methods (gcFront and gcSwarms), we used the largest library available to assess model performance over different search times. The selected time was determined by applying the elbow method over the generated data (**Supplementary Figure 5A-D**).

### 4 Analysis of ensemble model training process

To better understand the suitability of the data generated through gcFront and gcSwarms, as well as the performance of the generated ensemble models, several analyses were performed. Those analyses not only concern the diversity of the generated libraries of designs, but also the predictive performance of the generated models using those libraries or previously unseen data.

#### 4.1 Comparative analysis of training datasets

As is known in the field, an ML model performs as well as the data quality used for its training. In this regard, important needs to be fulfilled are the size and diversity of both the feature variables and the target variable to learn.

Libraries are assessed for their suitability according to its size and diversity. Regarding the size, the total number of unique designs meeting SD criteria was used. Regarding the characterization of their diversity, one frequency-based and other distance-based metric were implemented:

**Shannon Entropy per position**. This frequency-based metric was calculated using the GC design as a binary vector according to (*Def*. 1). Being *t* the number of candidate reactions computed by the SD pipeline, and *d* being the maximum number of KOs. Each vector position *i* is considered as a binary variable for whose Shannon Entropy is computed according to (8), being pi the proportion of 1s at position *i*. After that, the metric describing the diversity of the library is computed as the average of all variable entropies.

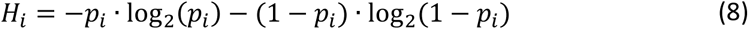

**Median Pairwise Jaccard Distance**. This distance-based metric was computed using the pdist function of the scipy python package^65^. This function computes every pairwise distance between all designs by calculating the difference between 1 and the JI computed following (1). The median of those values is used as the final diversity metric. Detailed implementation of both can be found in the file *“Method_comparation.ipynb”* present in the repository.

#### 4.2 Model performance with synthetic data

When training dataset size and complexity varies significantly across methodologies, standard performance parameters may not properly address the comparation of ML model performances. To tackle this comparation problem, we compute representative design spaces applying the DOE method LHS for all bioprocesses where designs are found using both SD algorithms. For generating such a design population, we will sample with LHS a different number of designs depending on the number of deletions (*d)* it contains, considering (9).

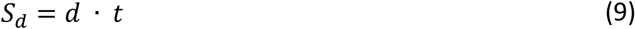

Being *Sd* the sample size for a design of order *d* within a design space of *t* reaction targets. With this approach, we will cover all the first order designs and obtain a representative sample for the rest of them, the size of it proportional to the design order. In this way we not only have a design space representation containing all design orders but also containing more examples to learn for the cases with a higher number of potential combinations. This generates synthetic designs that are more representative of the real variability of each design space in comparison with the training datasets, favoring a fair comparation.

Once the synthetic design space is generated, a residual analysis is performed to estimate the SD methodology performance on model generation. For this, true values of the scoring variable are computed by using the scoring function for it present in “*DesignScores.py*” file, within the *utils* folder of the ML4SD GitHub repository. On the other hand, predictions are made using bioprocess-specific ensemble models, from which SEs for a given true value of the variable are calculated (**Supplementary Figure 6**). Finally, for the overall estimation of predictive performance, the range-normalized root mean squared error (NRMSE) is calculated for each bioprocess-specific ensemble model (**A**).

### 5 Integration of ML4SD in a DBTL for strain-design

To generate the computational framework for implementing a DBTL cycle, we have chosen the *jupyter notebook* platform, as it displays a user-friendly interface while maintaining the capability of directly editing relevant variables affecting the workflow^66^. In our case, those variables are parameters affecting the strain-design algorithm (gcSwarms), the active-learning algorithm or the redirection of workflow output, whose values selection is explained below.

#### 5.1 Generation of ML-guided recommendations

The recommendations made are used as training data informing our ensemble model in following cycles, so capturing the model response to each feature within the design space is crucial to guide our DBTL cycle search. Considering this, the best option will be to simulate the entire design step and predict the performance for each design to later compute the individual deletion contribution to the final score. However, this design space is enormous, even for a small set of target reactions and therefore its computation is impractical (2).

To tackle this problem, we compute a representative design space using the DOE method LHS, following the approach described previously (section of 4.2 MM), which favors the representation of all features variability. Once we have computed a set of designs that is representative of the design space, the next step is to compute the feature-wise contribution to the score. To do so, we adapted the methodology described in a previous work concerning ML-guided recommendations^56^.

Specifically, we use a trained model to predict the performance of all designs in the computed set. Next, a threshold is introduced between 0 and the maximum predicted score. Within this range, a fixed number of evaluations are made in which we compute the frequency of each deletion, considering only the designs above the selected threshold for each step. In this way, as the threshold is increased, the statistical contribution of each deletion to the remaining part of the design space can be followed, expecting deletions with no significant contribution to higher scores to be underrepresented. Finally, to capture the relative importance of each deletion from the previous step to the target score, we compute the area under the curve (*AUC*) of the above-mentioned plot for each deletion. Each value is normalized so the sum of all AUCs gives one, representing our ML-based sampling probability distribution for a given score. With those steps, and according to (3) we end up generating *S(*D), that we will use to generate recommendations thereafter ().

#### 5.2 Feature importance analysis along DBTL cycles to elucidate large effects-deletions

This section describes the explainable ML methodology used for a better understanding of the decision-making process of the models. For this study, feature importance was calculated using the *Explainer* class of the *SHAP* python package^67^, as this is the most appropriate way for an ensemble model. The analysis was performed over each cycle model, thus registering the learning process through all the DBTL.

Firstly, Shapley values are computed for the best percentile of the cycle library. For this, we use the cycle-specific model and a representation of the training dataset given by 100 *kmedoids* extracted from the design library with *sklearn* python package^59^. Next, we get only the importance values for deletions (when a feature has a value of 1) and calculate its importance and presence. The first parameter is computed as the mean Shapley value across all explained designs, normalized by the maximum mean absolute Shapley value; while the second is the percentage of designs (considering the selected percentile) in which a deletion is present. For a detailed view of the analysis please see the “*SHAP_explainability.ipynb*” file.

### 6 Computational resources

Being the present study purely computational, the materials used consists basically of different types of computer networks available in the work center along with my personal computer. The technical details of each of these can be found in ***Table 4***.

**Table 4.** Hardware specifications of computer systems used in this work.

| *System Name* | *Type* | *OS* | *CPUs* | *RAM* |
| --- | --- | --- | --- | --- |
| *Modern-14-B11MO* | PC | Ubuntu 20.04.6 LTS | 8 | 15.3 GB |
| *Elbrus* | HPC | Ubuntu 20.04.4 LTS | 104 | 1.48 TB |

## Supplementary figures

**Supplementary Figure 1.**
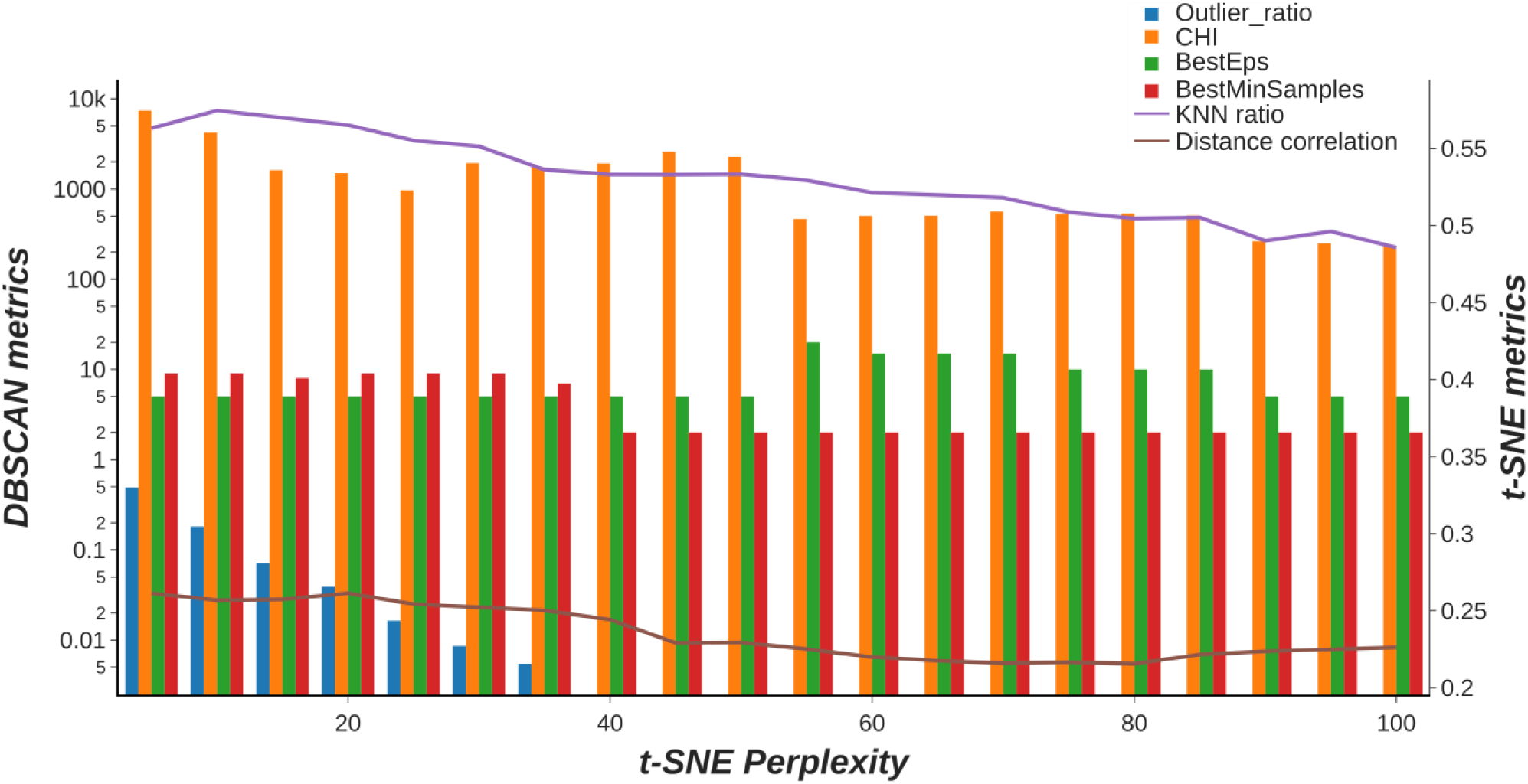
Setup and quality metrics for selecting KT chemical space. The plot shows metrics concerning dimensionality reduction and clustering setup for generation the chemical space of iJN1462 metabolites. The quality of the dimensionality reduction performed by t-SNE was assessed through the ratio of K-nearest neighbors (light purple line). Concerning the clustering through DBSCAN, the plot shows the hyperparameter setup (bars) and its quality assessment through the Calinski–Harabasz index (CHI) and outlier ratio.

**Supplementary Figure 2.**
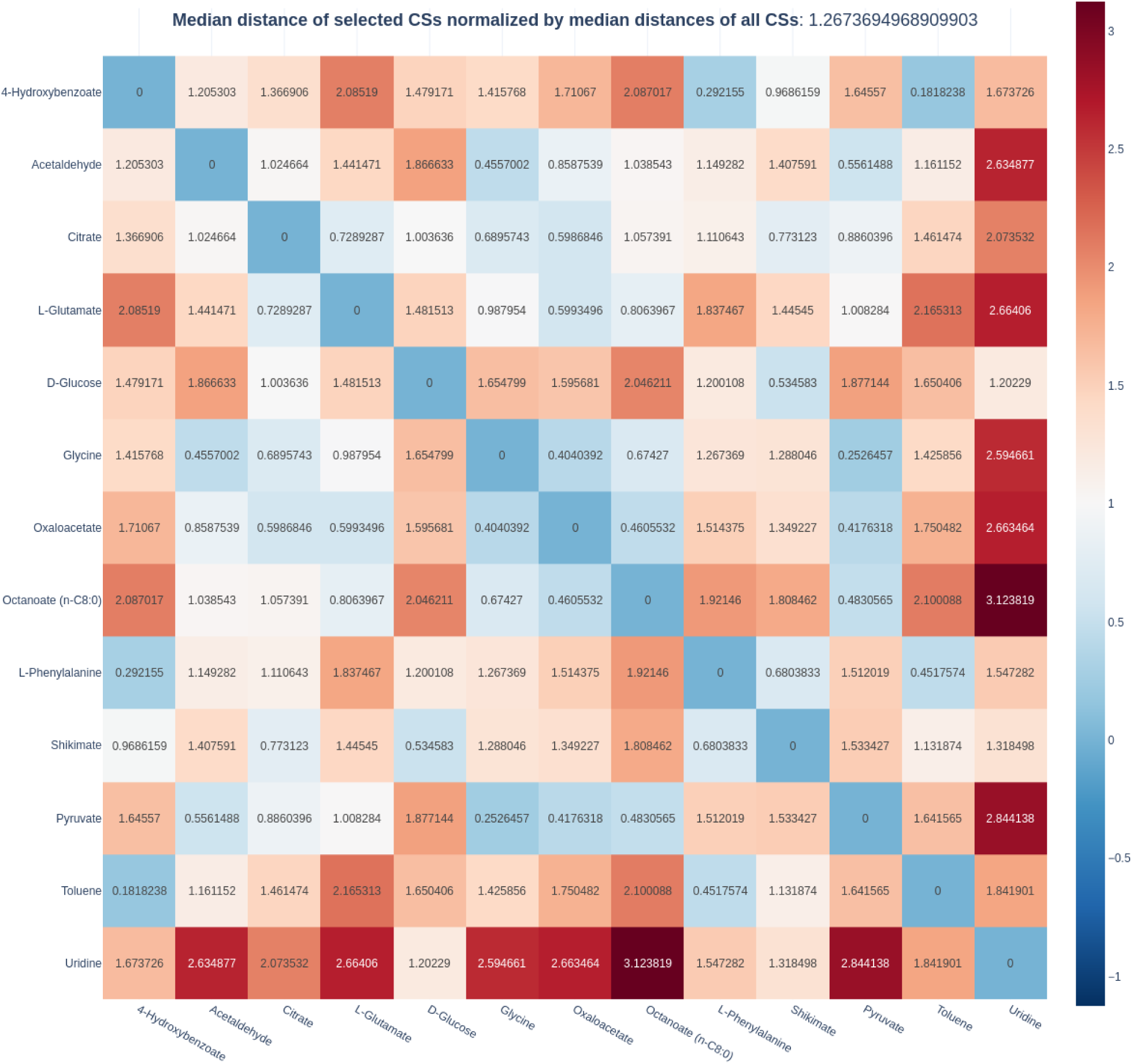
Relative distances of carbon sources in iJN1462. Heatmap representing pairwise distances of selected carbon sources normalized by the median pairwise distances of all carbon sources in *iJN1462*. Distance in this plot refers to Euclidean distances between a pair of metabolites within the chemical space represented through t-SNE.

**Supplementary Figure 3.**
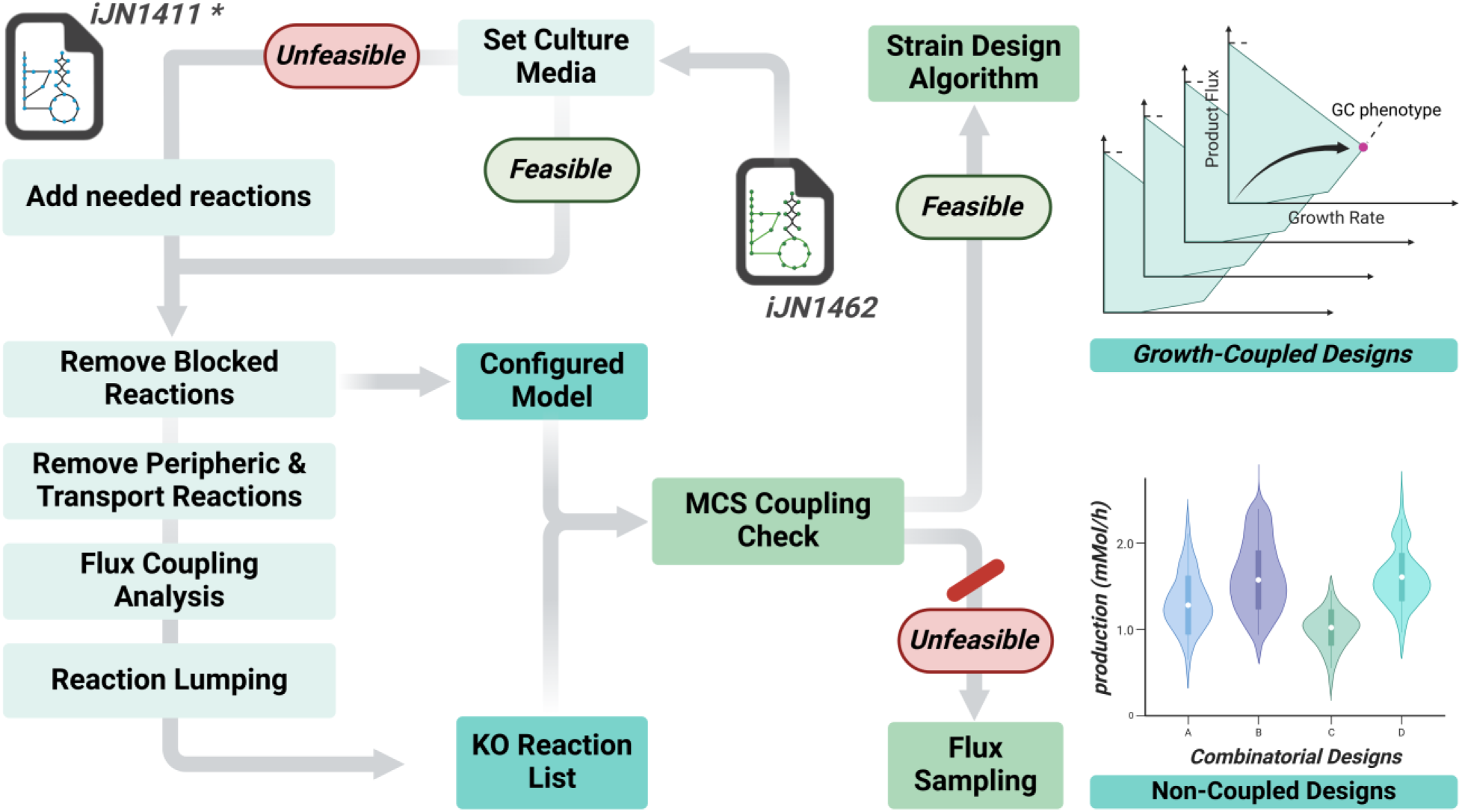
In-house strain design pipeline for library generation. The pipeline starts with a base GEM (*iJN1462*) that, if necessary, is expanded using a repository model containing all necessary reactions. In this study, such a repository model is selected from a previous study (*iJN1411\**)^39^. After this, a reduction step is performed that removes from the GEM non-target biomass reactions and reactions blocked in the bioprocess condition. From these reactions of the configured model, the final Knock-Out (KO) candidate list is generated by excluding essential, transport, and non-GPR reactions, followed by a user-set carbon limit filtering and reaction lumping. With both configured model and candidate KOs, a Minimal Cut Sets (MCS) analysis is performed to pre-filter non-feasible growth coupled bioprocesses. If a bioprocess pass the test, a strain design search algorithm is run to provide the growth-coupled designs.

**Supplementary Figure 4.**
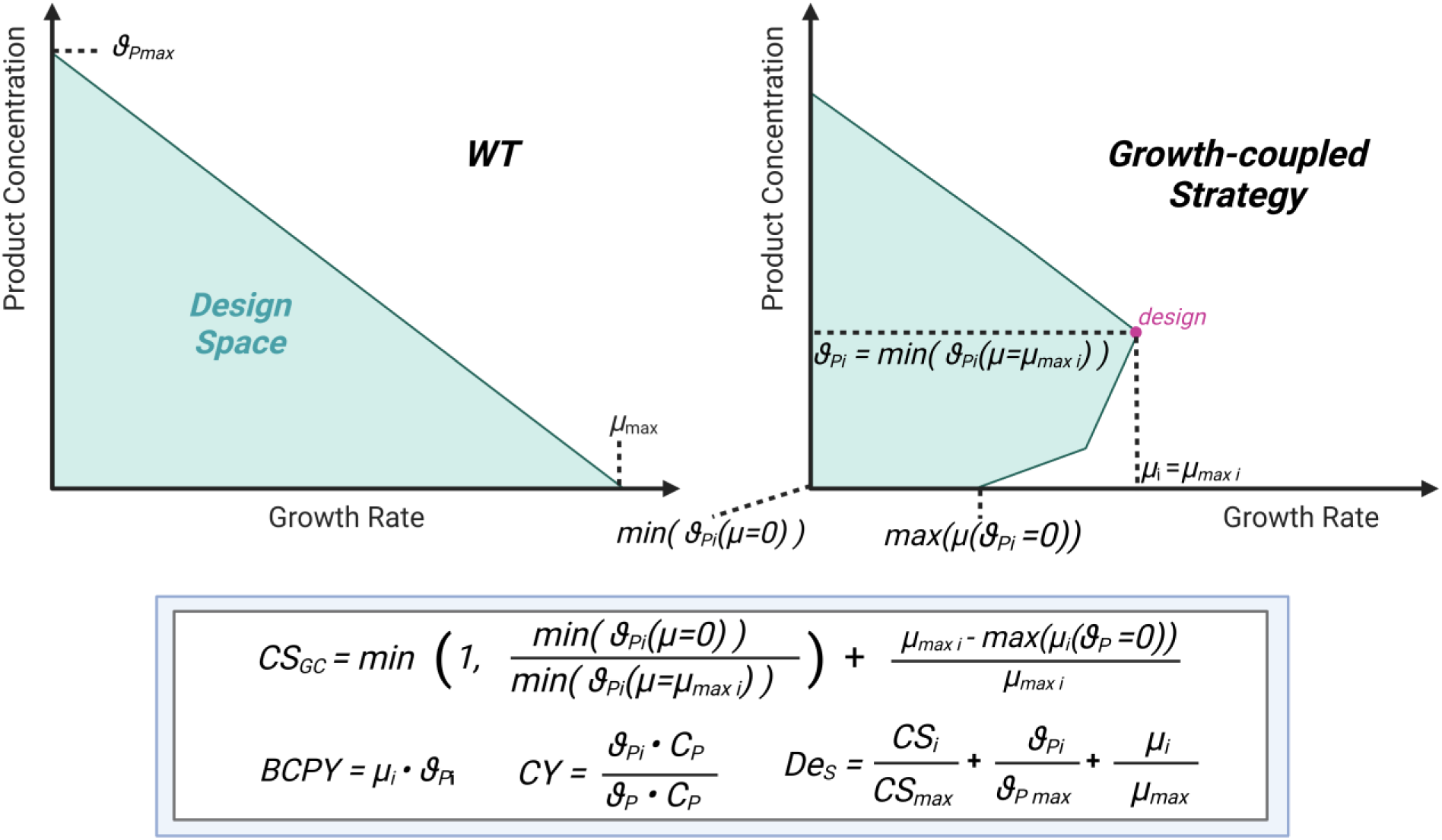
Graphic explanation of scores used in gcSwarms. Parameters of the equations described within the paper body are graphically represented within a production envelope plot. This plot represents the GEM feasible space for a given bioprocess, which in the *WT* usually consists of a tradeoff between the product titter and the growth rate of the selected microbial chassis.

**Supplementary Figure 5.**
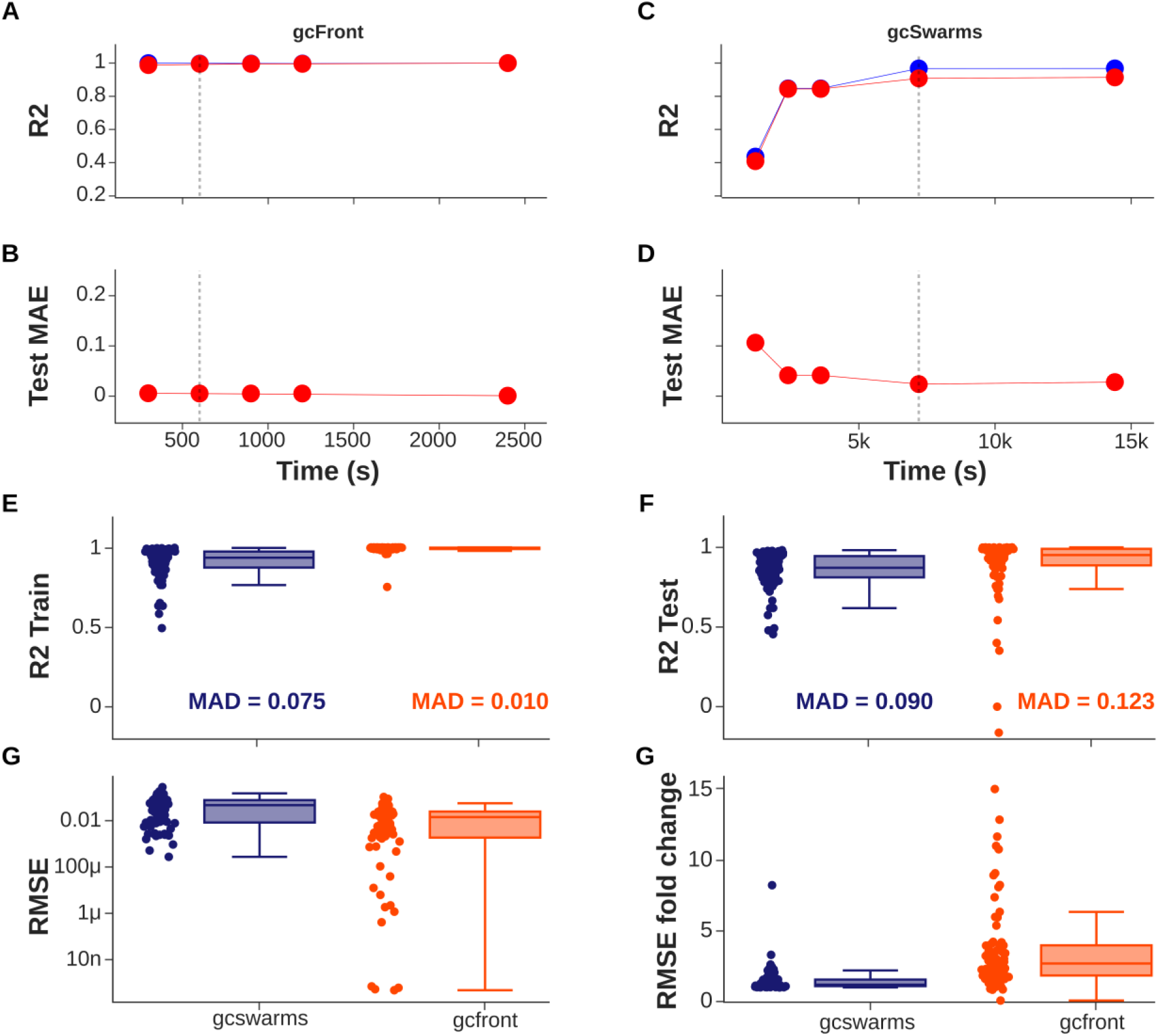
Ensemble models evaluation over train and test dataset. (**A**, **C**) Scatter plots that show the dependence of the search time used for *Autosklearn* to find a proper ensemble model and its R^2^ for train (blue) and test (red) datasets belonging to gcFront (**A**) or gcSwarms (**B**). (**B**, **D**) Scatter plots that show the dependence of the search time used for *Autosklearn* to find a proper ensemble model and its mean absolute error (MAE) for the test dataset belonging to gcFront (**B**) or gcSwarms (**D**). (**E**, **F**) Bar plots showing the predicted power (as R^2^) of all generated ensemble models over the train (**E**) and test (**F**) datasets for both strain design algorithms used in this paper. (**G**) Bar plots of root of mean squared errors (RMSE) of the ensemble models over the test dataset. (**H**) Bar plots with fold change of RMSE values of the generated ensemble between the train and test datasets. From **E** to **H** dots represent values of each of the ensemble models.

**Supplementary Figure 6.**
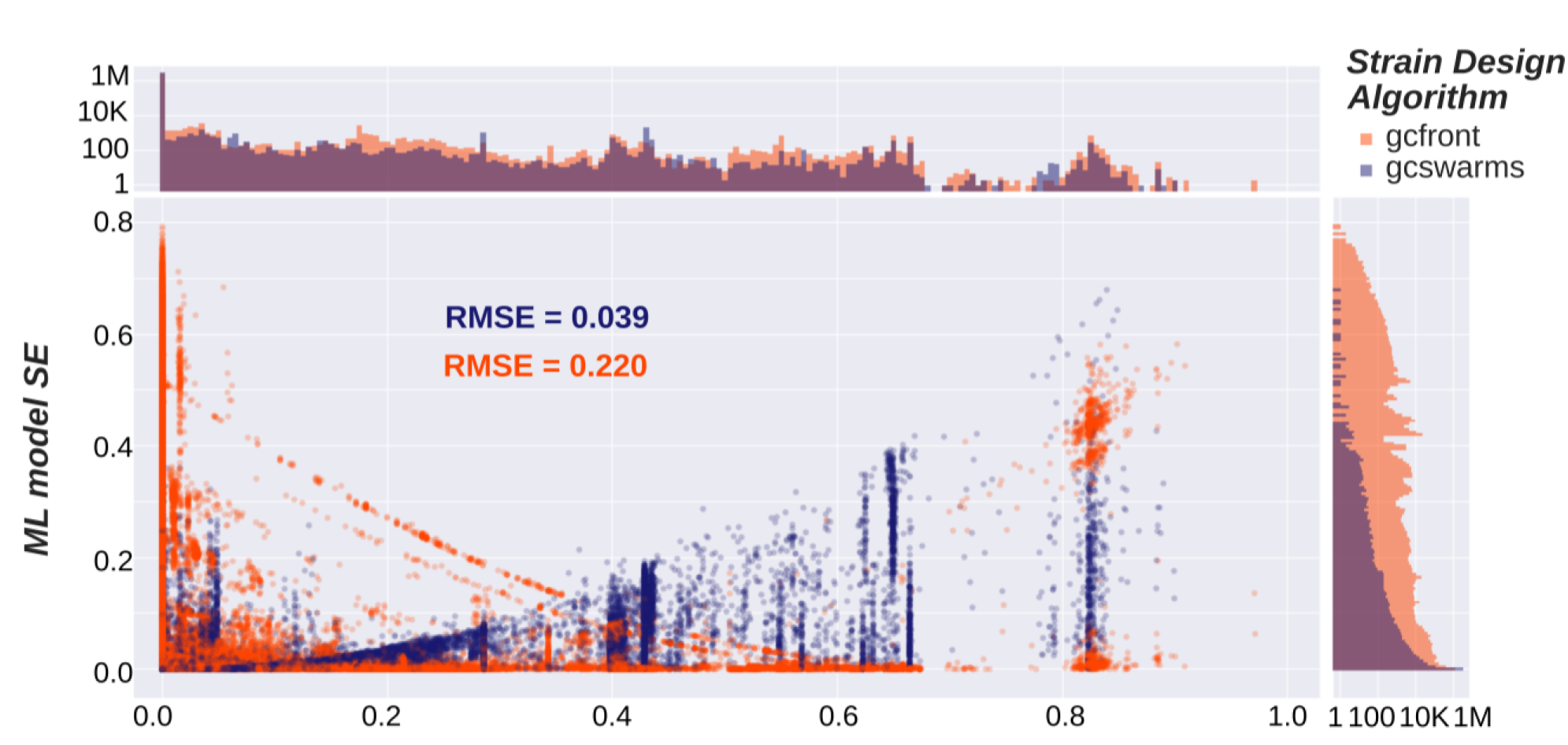
Aggregated squared errors from ensemble models predictions of synthetic data. Composed plot built from synthetic data evaluation. Data corresponds to ensemble models trained with design libraries of bioprocesses shared between gcFront and gcSwarms. The scatter plot represents the relationship between the true value of design performance according to GEM validation and the squared error of model prediction. Dots correspond to prediction and evaluation data of unique designs. Histograms at the top and the right of the figure show the distribution of true GEM values and SEs of ensemble models respectively. Note that both histograms have logarithmic axis for better visualization.

**Supplementary Figure 7.**
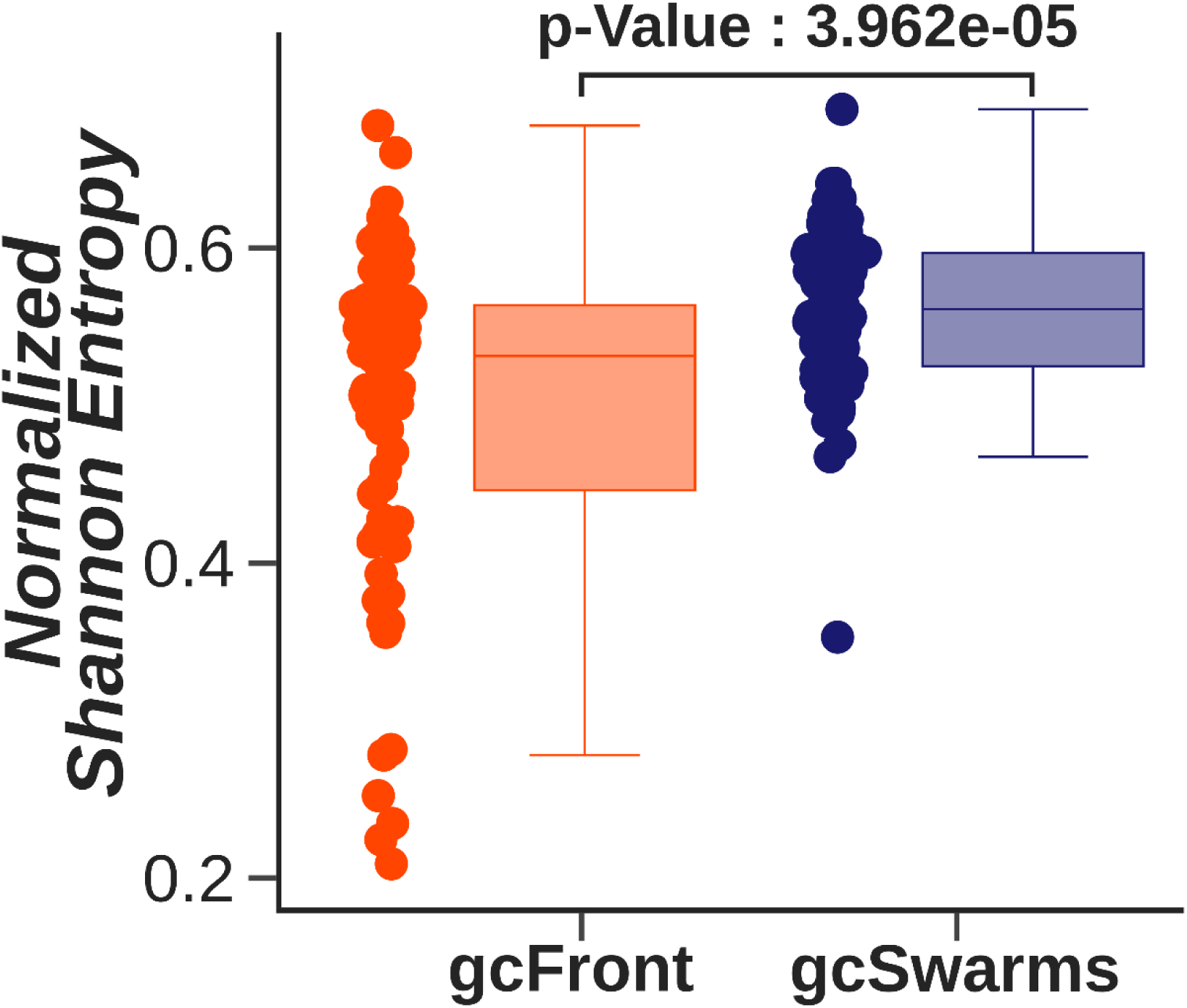
Entropy of initial strain design libraries. Bar plot showing normalized Shannon entropy values obtained for each of the GC libraries. This metric was computed by treating designs as binary vectors, using each position in it as a unique binary variable*(8)*. The final value is the variable-normalized sum of all those variables.

**Supplementary Figure 8.**
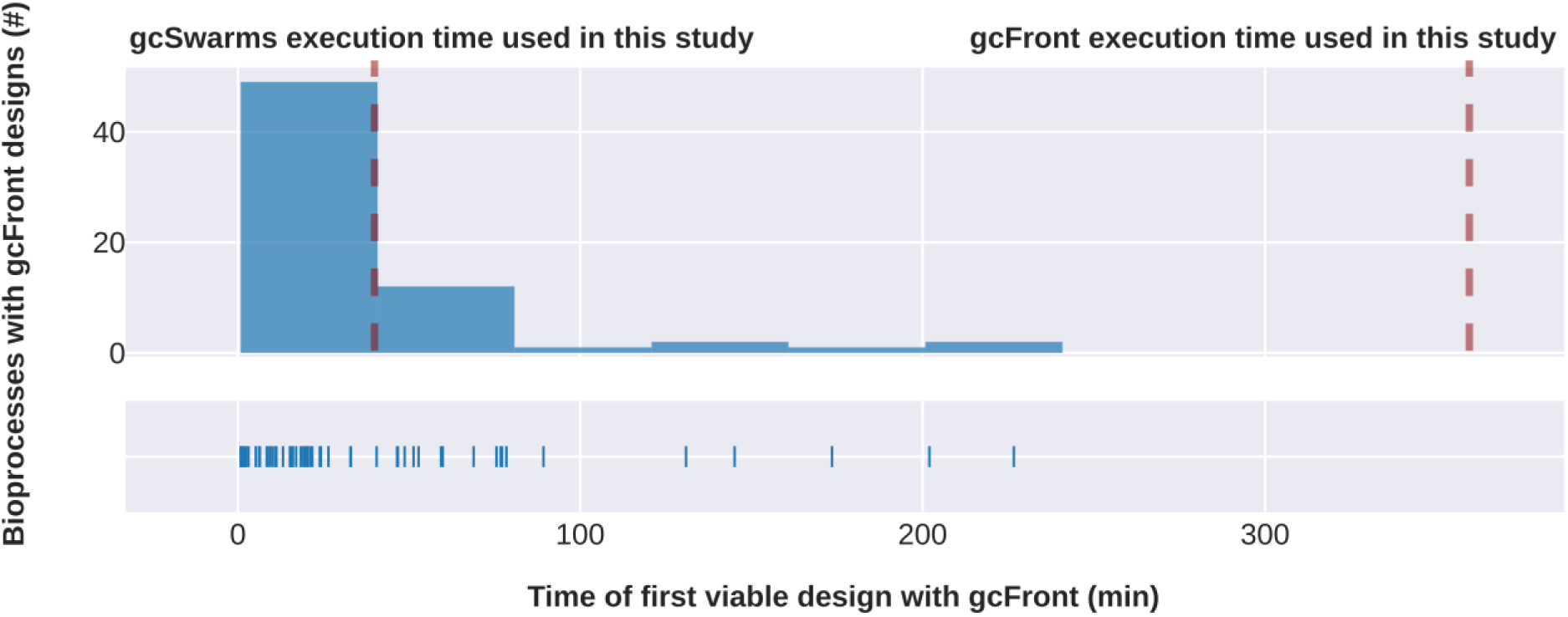
Distribution of the time gcFront needs to find the first viable design. Histogram plot where the distribution of times that gcFront needed to find the first design meeting with the criteria of this study are represented. Below, the same data is represented through individual lines. Also, red dashed lines are represented that represent gcSwarms (left) and gcFront (right) total execution time in minutes.

**Supplementary Figure 9.**
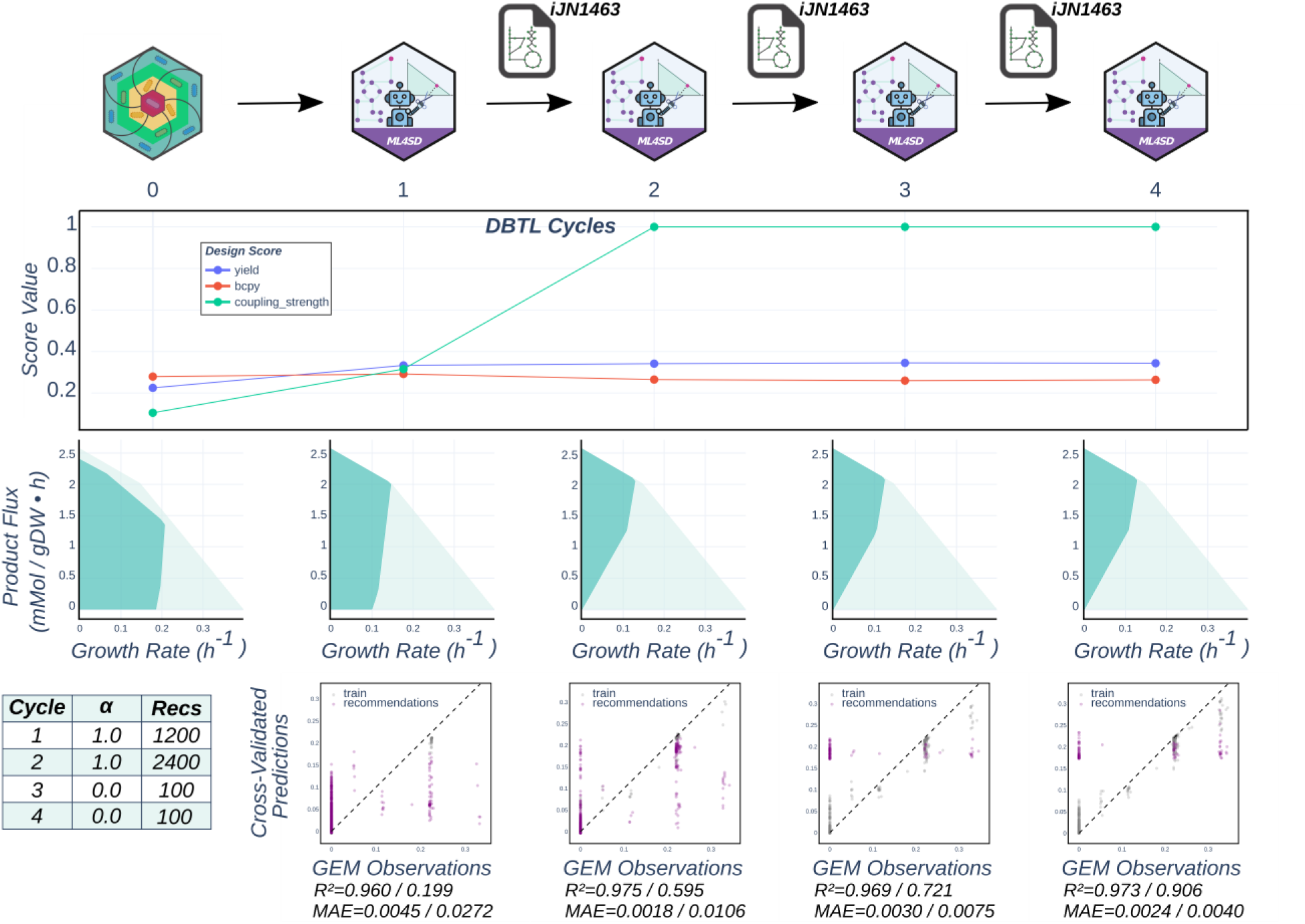
Results of the replicate 1 from ML4SD campaign for CY. Progression of fitness scores of best designs across DBTL rounds are shown (top). The dots and lines represent the first replicate of *CY* campaign. Below, the envelopes of the best design (dark blue) for each round are represented in comparison with the *WT* (light blue). At the bottom, a summary table recapitulates the DBTL parameter setup chosen for the present case study. Following this, predicted cross-validations are plotted against true values according to GEM simulations for each DBTL round. Below each graph R^2^ and MAE values are shown. Those correspond to ensemble model predictions over the library used as training dataset (left) and over predictions of the library after all round recommendations have been added (right).

**Supplementary Figure 10.**
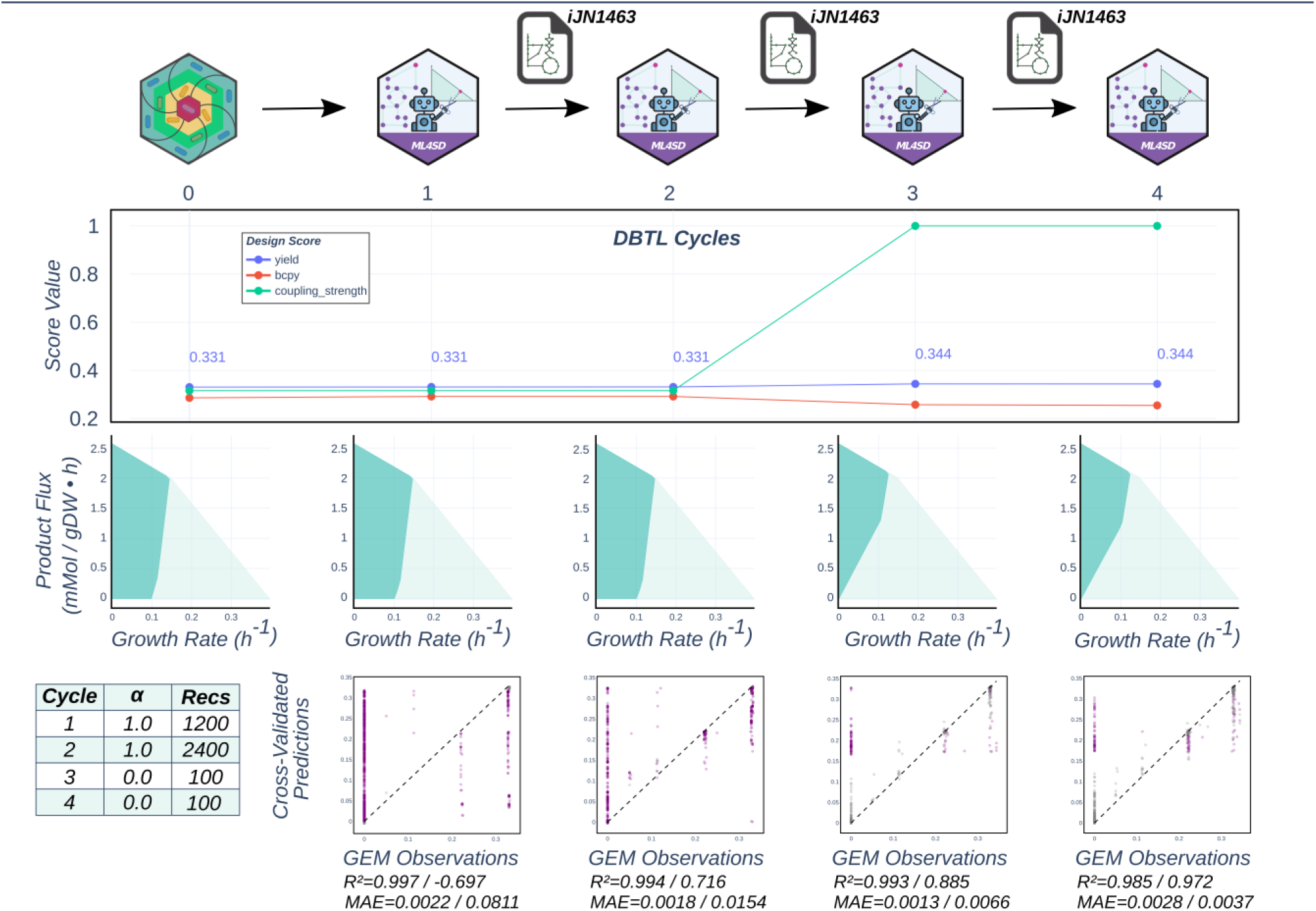
Results of the replicate 3 from ML4SD campaign for CY. Progression of fitness scores of best designs across DBTL rounds are shown (top). The dots and lines represent the third replicate of *CY* campaign. Below, the envelopes of the best design (dark blue) for each round are represented in comparison with the *WT* (light blue). At the bottom, a summary table recapitulates the DBTL parameter setup chosen for the present case study. Following this, predicted cross-validations are plotted against true values according to GEM simulations for each DBTL round. Below each graph R^2^ and MAE values are shown. Those correspond to ensemble model predictions over the library used as training dataset (left) and over predictions of the library after all round recommendations have been added (right).

**Supplementary Figure 11.**
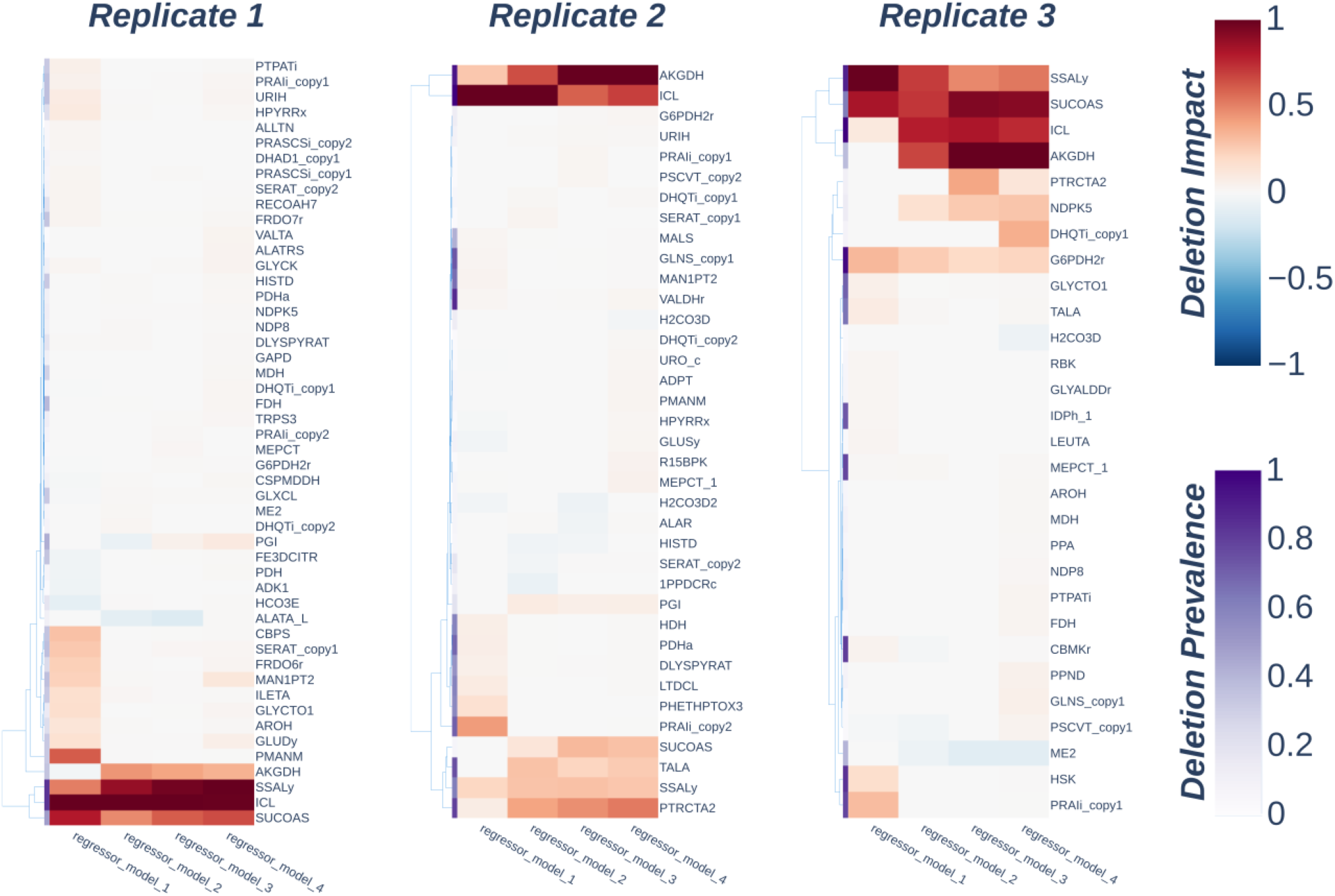
Deletion importance values extracted from SHAP analysis of ensemble models of CY campaign. Clustered heatmaps recapitulating importance and prevalence of deletions according to ensemble models of each DBTL round for all replicates of *CY* campaign. The first parameter is computed from Shapley values of the top percentile designs within the generated library, while the second is the percentage of those designs in which a deletion is present (see MM section 5.2).

**Supplementary Figure 12.**
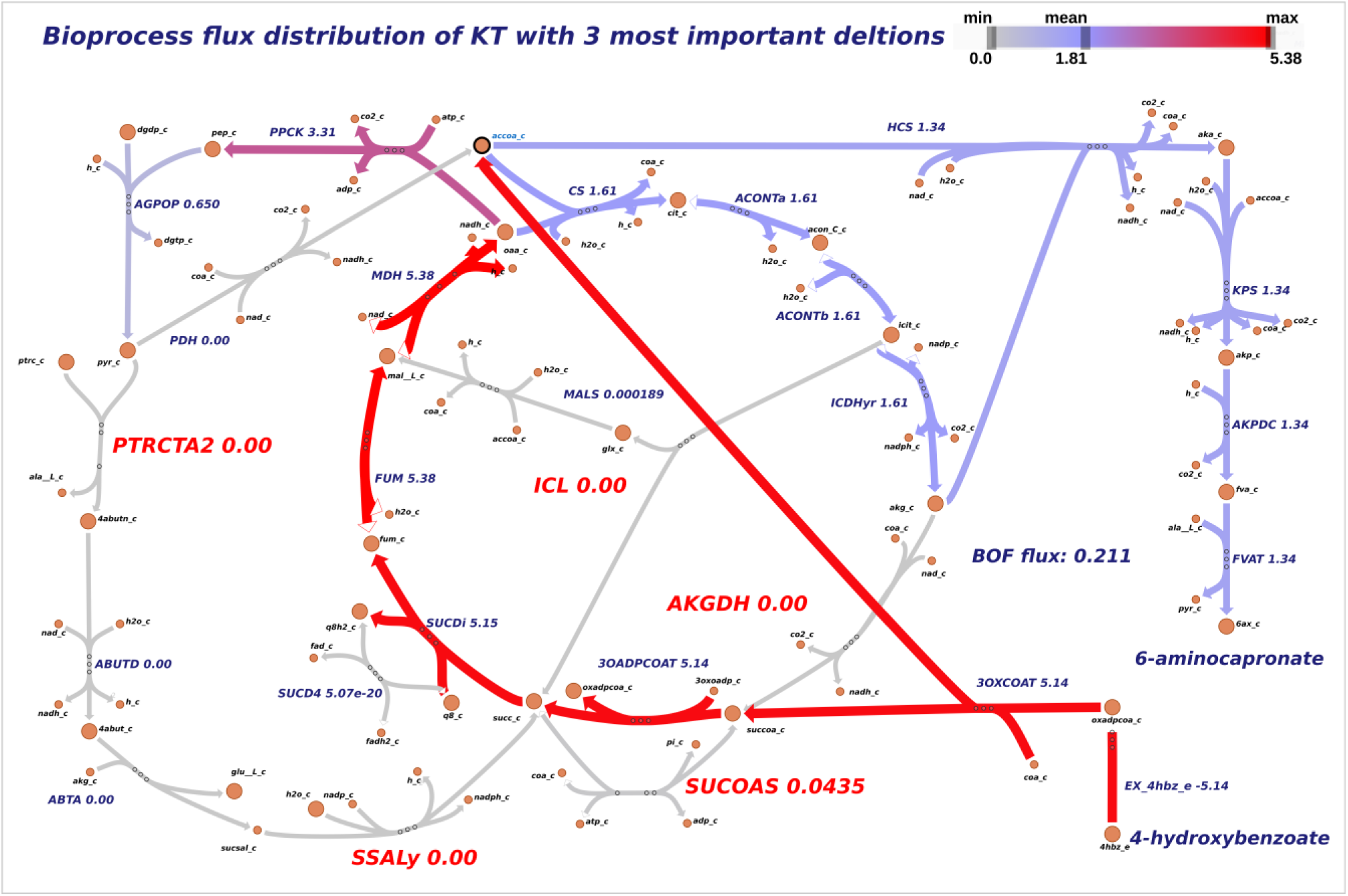
Metabolic flux map showing the effects of the minimal design. Metabolic map generated by using *escher* online application and visualized through its python package^68^. The map shows the flux distribution that the configured model predicts when we applied the top 3 deletions according to ensemble models (*AKGDH*, *ICL* and *PTRCTA* / *SSALy*). Top 5 reactions along all ensemble models are highlighted in red color.

**Supplementary Figure 13.**
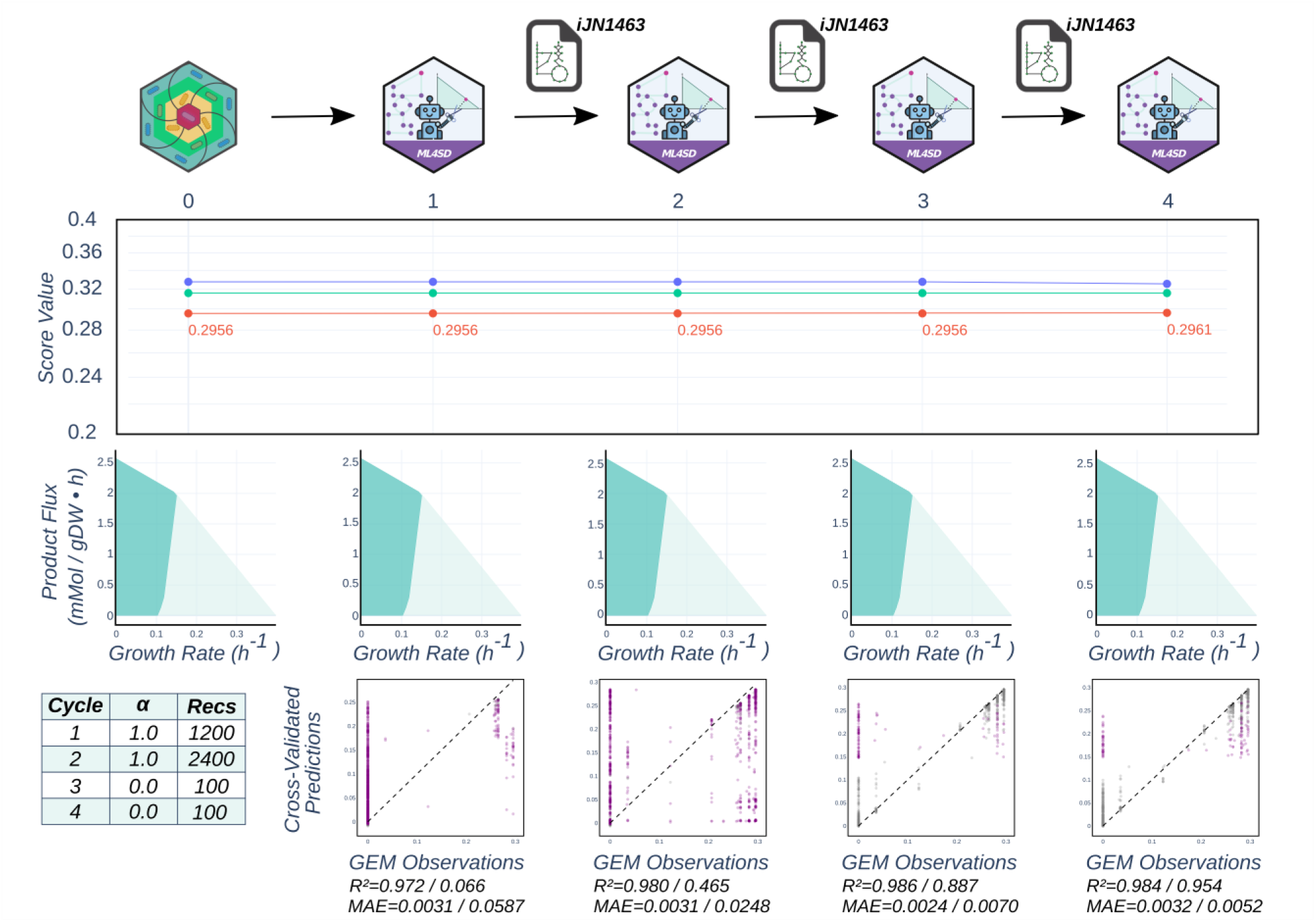
Results of replicate 1 from DBTL campaign optimizing design BCPY. Progression of fitness scores of best designs across DBTL rounds are shown (top). The dots and lines represent the first replicate of *BCPY* campaign. Below, the envelopes of the best design (dark blue) for each round are represented in comparison with the *WT* (light blue). At the bottom, a summary table recapitulates the DBTL parameter setup chosen for the present case study. Following this, predicted cross-validations are plotted against true values according to GEM simulations for each DBTL round. Below each graph R^2^ and MAE values are shown. Those correspond to ensemble model predictions over the library used as training dataset (left) and over predictions of the library after all round recommendations have been added (right).

**Supplementary Figure 14.**
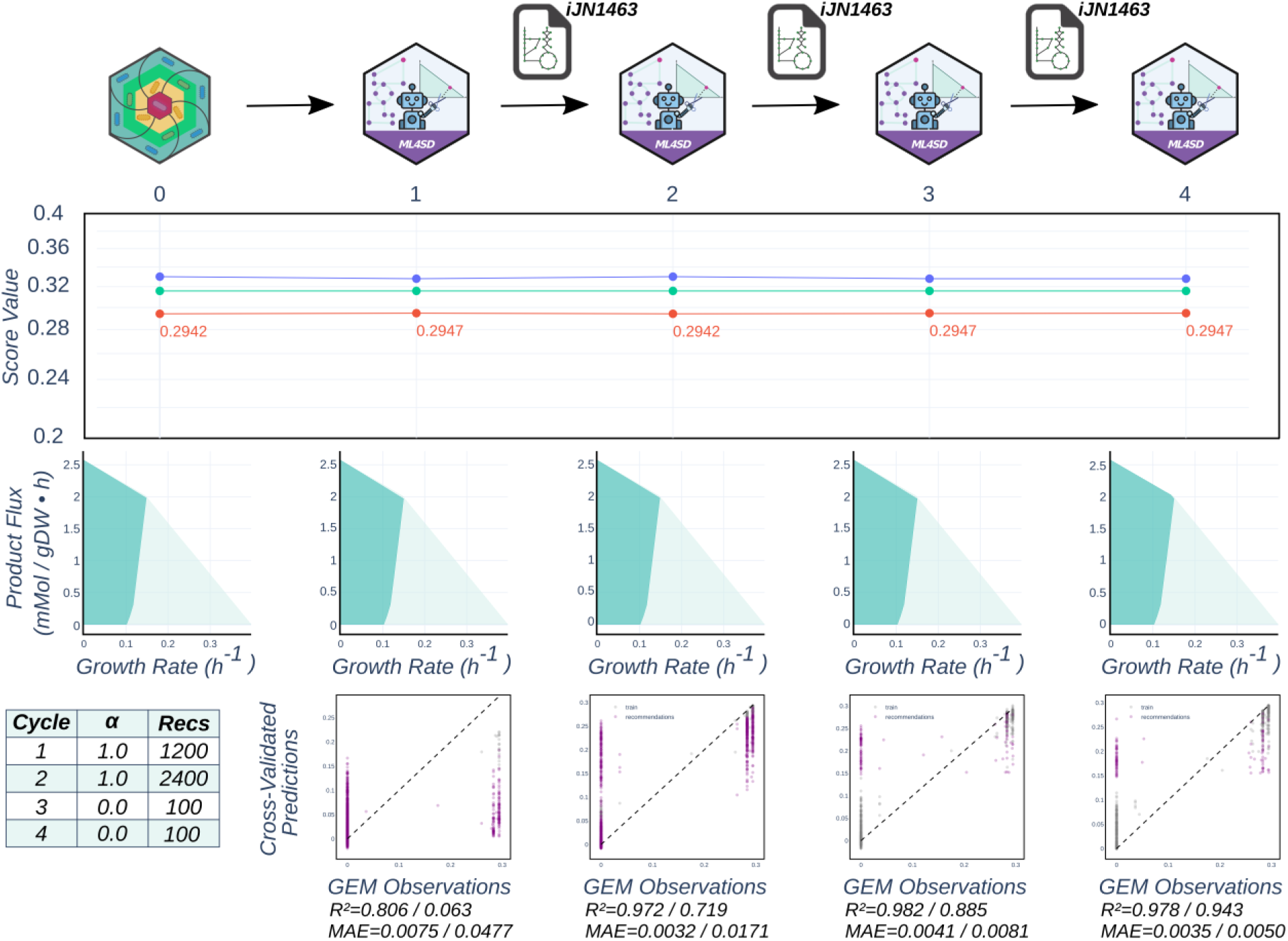
Results of replicate 2 from DBTL campaign optimizing design BCPY. Progression of fitness scores of best designs across DBTL rounds are shown (top). The dots and lines represent the second replicate of *BCPY* campaign. Below, the envelopes of the best design (dark blue) for each round are represented in comparison with the *WT* (light blue). At the bottom, a summary table recapitulates the DBTL parameter setup chosen for the present case study. Following this, predicted cross-validations are plotted against true values according to GEM simulations for each DBTL round. Below each graph R^2^ and MAE values are shown. Those correspond to ensemble model predictions over the library used as training dataset (left) and over predictions of the library after all round recommendations have been added (right).

**Supplementary Figure 15.**
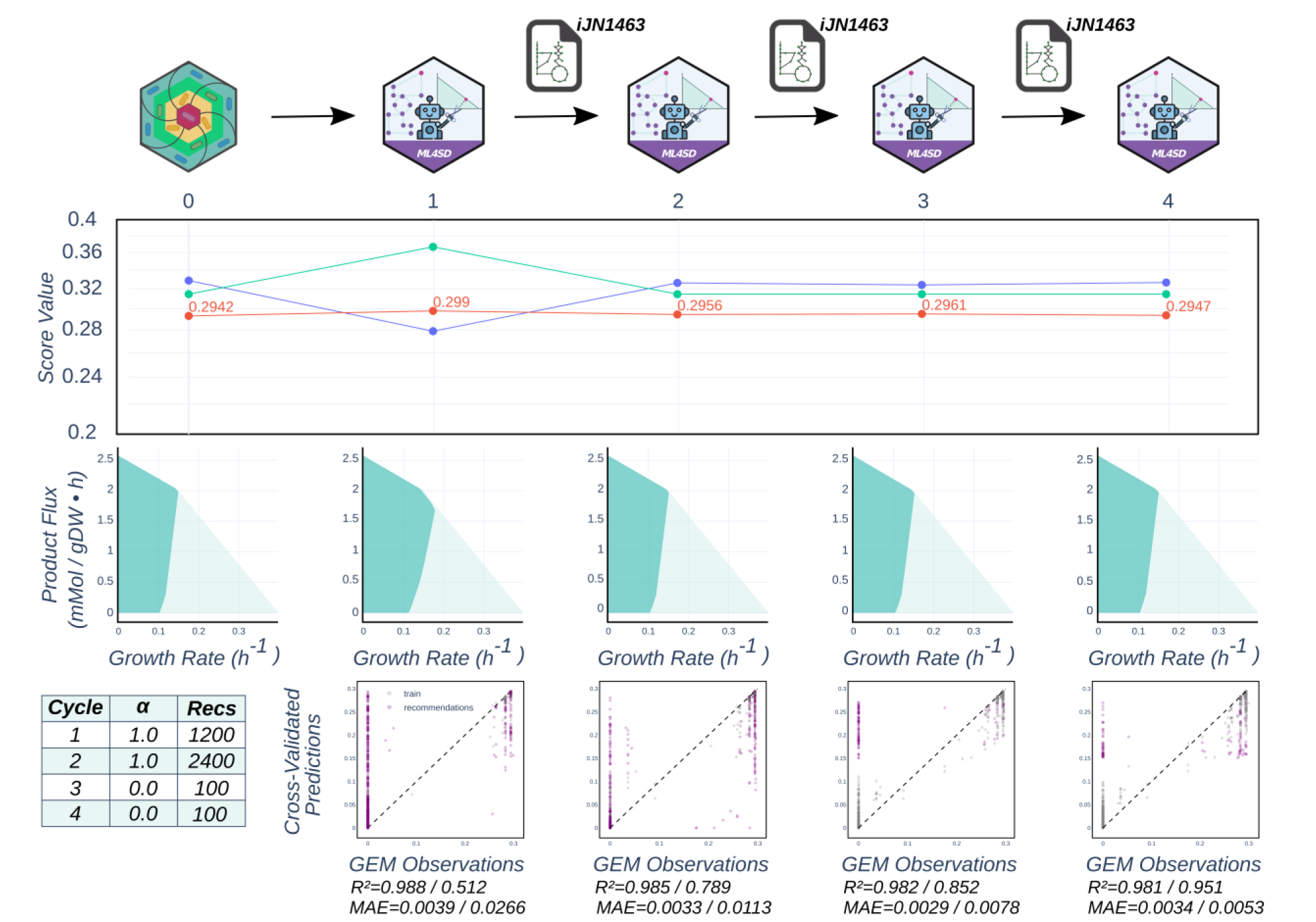
Results of replicate 3 from DBTL campaign optimizing design BCPY. Progression of fitness scores of best designs across DBTL rounds are shown (top). The dots and lines represent the third replicate of *BCPY* campaign. Below, the envelopes of the best design (dark blue) for each round are represented in comparison with the *WT* (light blue). At the bottom, a summary table recapitulates the DBTL parameter setup chosen for the present case study. Following this, predicted cross-validations are plotted against true values according to GEM simulations for each DBTL round. Below each graph R^2^ and MAE values are shown. Those correspond to ensemble model predictions over the library used as training dataset (left) and over predictions of the library after all round recommendations have been added (right).

**Supplementary Figure 16.**
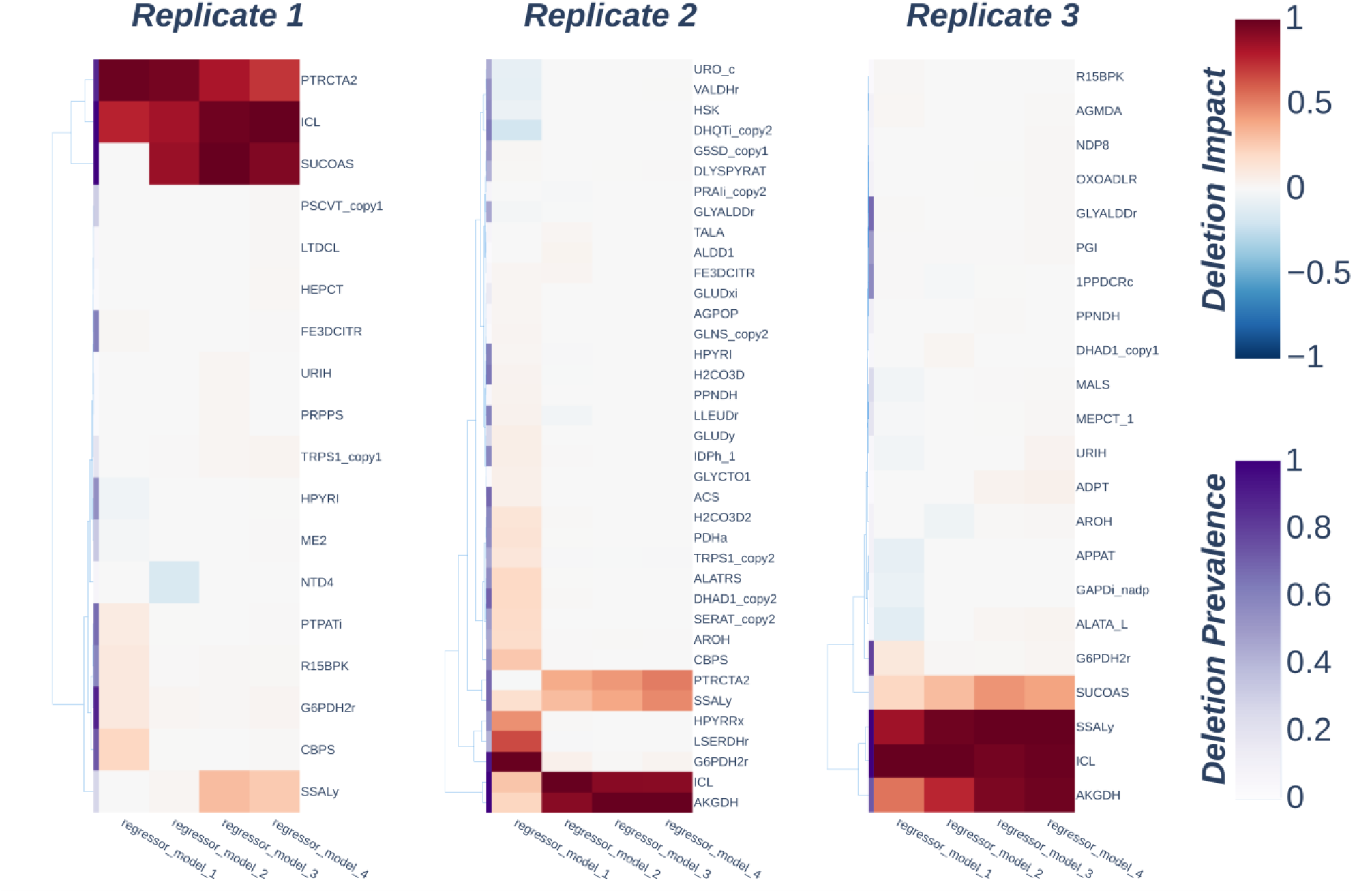
Deletion importance values extracted from SHAP analysis of ensemble models of BCPY campaign. Clustered heatmaps recapitulating importance and prevalence of deletions according to ensemble models of each DBTL round for all replicates of *BCPY* campaign. The first parameter is computed from Shapley values of the top percentile designs within the generated library, while the second is the percentage of those designs in which a deletion is present (see MM section 5.2).

**Supplementary Figure 17.**
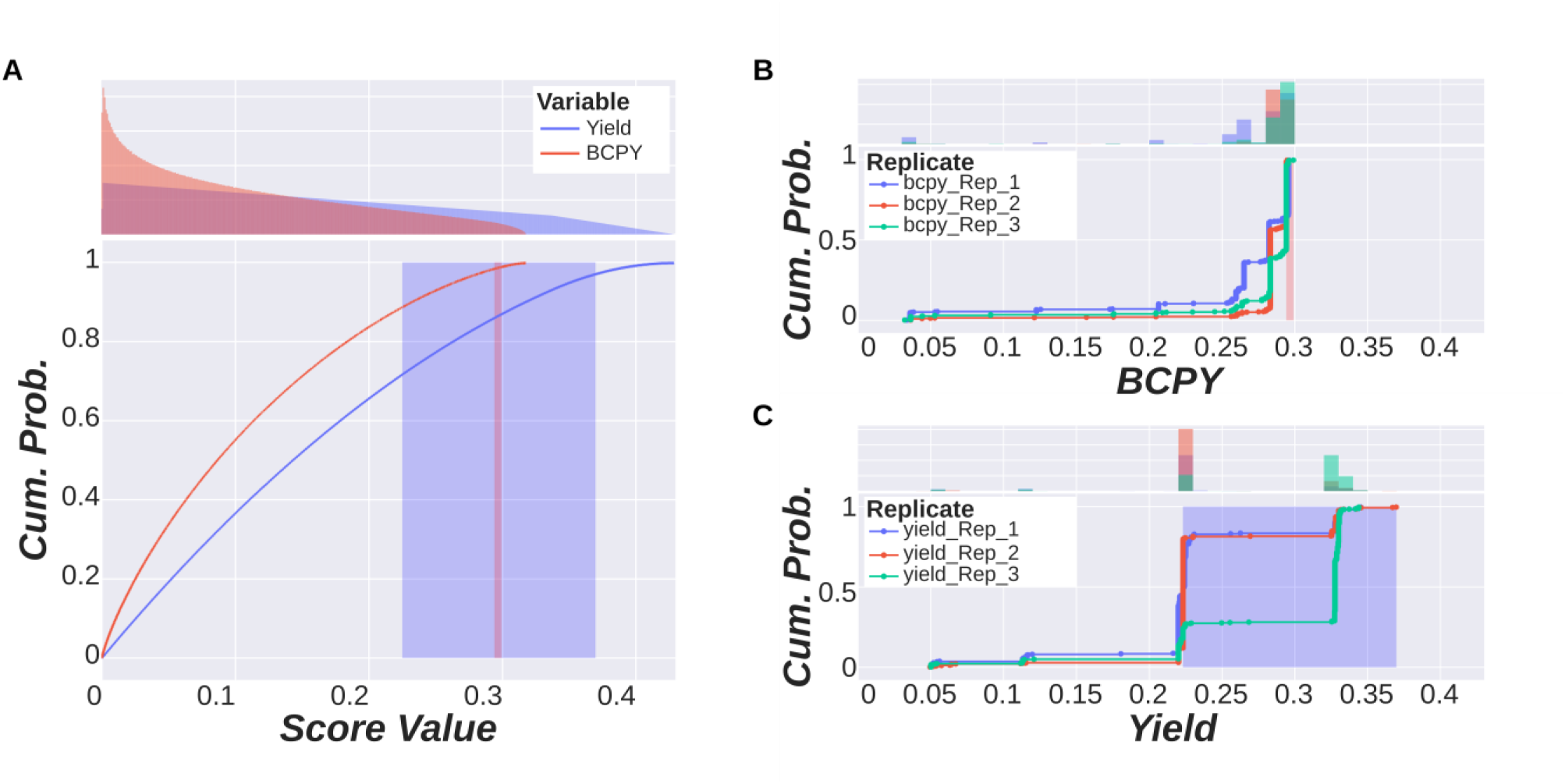
Theoretical and empirical cumulative probability of CY and BCPY. Cumulative distribution plots of performance scores used in this study. The plot considering theoretical distribution is made by assuming that all points within the production envelope are equally probable (**A**). Also, plots are made for empirical cumulative probability considering only the viable designs found among 3 ML4SD replicates concerning *BCPY* (**B**) and *CY* (**C**) campaigns. The colored area in each plot represents the exploitative range, which is the difference between the best design score of *gcSwarms* and the best design found in the final library. Above each plot, a histogram counts the number of designs across different score values.

## Notes

https://github.com/extrevaro/ML4SD

### Competing Interest Statement

The authors have declared no competing interest.

## References

1. Pavan, M. et al. Advances in systems metabolic engineering of autotrophic carbon oxide-fixing biocatalysts towards a circular economy. Metab. Eng. 71, 117–141 (2022).

2. Andhalkar, V. V. et al. Integrated Biorefinery Design with Techno-Economic and Life Cycle Assessment Tools in Polyhydroxyalkanoates Processing. Macromol. Mater. Eng. n/a, 2300100.

3. An automated Design-Build-Test-Learn pipeline for enhanced microbial production of fine chemicals | Communications Biology. https://www.nature.com/articles/s42003-018-0076-9.

4. Asin-Garcia, E., Fawcett, J. D., Batianis, C. & Martins dos Santos, V. A. P. A snapshot of biomanufacturing and the need for enabling research infrastructure. Trends Biotechnol. https://doi.org/10.1016/j.tibtech.2024.10.014 (2024) doi:10.1016/j.tibtech.2024.10.014.

5. A pipeline for the reconstruction and evaluation of context-specific human metabolic models at a large-scale | PLOS Computational Biology. https://journals.plos.org/ploscompbiol/article?id=10.1371/journal.pcbi.1009294.

6. Banerjee, D. et al. Addressing genome scale design tradeoffs in Pseudomonas putida for bioconversion of an aromatic carbon source. Npj Syst. Biol. Appl. 11, 1–13 (2025).

7. Maia, P., Rocha, M. & Rocha, I. In Silico Constraint-Based Strain Optimization Methods: the Quest for Optimal Cell Factories. Microbiol. Mol. Biol. Rev. 80, 45–67 (2015).

8. Banerjee, D. & Mukhopadhyay, A. Perspectives in Growth Production Trade-off in Microbial Bioproduction. RSC Sustain. https://doi.org/10.1039/D2SU00066K (2023) doi:10.1039/D2SU00066K.

9. Feist, A. M. et al. Model-driven evaluation of the production potential for growth-coupled products of Escherichia coli. Metab. Eng. 12, 173–186 (2010).

10. Legon, L., Corre, C., Bates, D. G. & Mannan, A. A. gcFront: a tool for determining a Pareto front of growth-coupled cell factory designs. Bioinformatics 38, 3657–3659 (2022).

11. Yang, Z. & Tamura, T. DBgDel: Database-Enhanced Gene Deletion Framework for Growth-Coupled Production in Genome-Scale Metabolic Models. Preprint at 10.48550/arXiv.2411.08077 (2025).

12. Lu, H., Xiao, L., Liao, W., Yan, X. & Nielsen, J. Cell factory design with advanced metabolic modelling empowered by artificial intelligence. Metab. Eng. 85, 61–72 (2024).

13. Zhang, J. et al. Combining mechanistic and machine learning models for predictive engineering and optimization of tryptophan metabolism. Nat. Commun. 11, 4880 (2020).

14. Carruthers, D. N. et al. Automation and machine learning drive rapid optimization of isoprenol production in Pseudomonas putida. Nat. Commun. 16, 11489 (2025).

15. Kugler, A. & Stensjö, K. Machine learning predicts system-wide metabolic flux control in cyanobacteria. Metab. Eng. 82, 171–182 (2024).

16. Tazza, G., Moro, F., Ruggeri, D., Teusink, B. & Vidács, L. MINN: A metabolic-informed neural network for integrating omics data into genome-scale metabolic modeling. Comput. Struct. Biotechnol. J. 27, 3609–3617 (2025).

17. Gotsmy, M. & Guillén-Gosálbez, G. Integrating Metabolic Networks into Hybrid Bioprocess Models. 2026.04.22.720062 Preprint at 10.64898/2026.04.22.720062 (2026).

18. Merzbacher, C. & Oyarzún, D. A. Applications of artificial intelligence and machine learning in dynamic pathway engineering. Biochem. Soc. Trans. 51, 1871–1879 (2023).

19. Kumar, P. et al. Active and machine learning-based approaches to rapidly enhance microbial chemical production. Metab. Eng. 67, 216–226 (2021).

20. Leavell, M. D., Singh, A. H. & Kaufmann-Malaga, B. B. High-throughput screening for improved microbial cell factories, perspective and promise. Curr. Opin. Biotechnol. 62, 22–28 (2020).

21. Yi, X. et al. Establishing a versatile toolkit of flux enhanced strains and cell extracts for pathway prototyping. Metab. Eng. 80, 241–253 (2023).

22. A Microfluidic Multiplex Sorter for Strain Development - Leal-Alves - Advanced Materials Technologies - Wiley Online Library. https://onlinelibrary.wiley.com/doi/10.1002/admt.202401209?msockid=2af65f865591648f293c4bb354e865e0.

23. Li, X. et al. Leveraging large language models for metabolic engineering design. Trends Biotechnol. https://doi.org/10.1016/j.tibtech.2026.03.026 (2026) doi:10.1016/j.tibtech.2026.03.026.

24. Cohn, D. A., Ghahramani, Z. & Jordan, M. I. Active Learning with Statistical Models. J. Artif. Intell. Res. 4, 129–145 (1996).

25. Borkowski, O. et al. Large scale active-learning-guided exploration for in vitro protein production optimization. Nat. Commun. 11, 1872 (2020).

26. Yang, Z. & Tamura, T. DeepGDel: Deep Learning-Based Gene Deletion Prediction Framework for Growth-Coupled Production in Genome-Scale Metabolic Models. IEEE Trans. Comput. Biol. Bioinforma. 22, 2252–2266 (2025).

27. Radivojević, T., Costello, Z., Workman, K. & Garcia Martin, H. A machine learning Automated Recommendation Tool for synthetic biology. Nat. Commun. 11, 4879 (2020).

28. Van Lent, P., Paz, S. M., Schmitz, J. & Abeel, T. Comparing metabolic engineering scenarios using simulated design-build-test-learn-cycles. Front. Bioeng. Biotechnol. 14, (2026).

29. Feurer, M. et al. Efficient and Robust Automated Machine Learning.

30. Noor, M. S. et al. Next-generation metabolic models informed by biomolecular simulations. Curr. Opin. Biotechnol. 92, 103259 (2025).

31. Sabzevari, M., Szedmak, S., Penttilä, M., Jouhten, P. & Rousu, J. Strain design optimization using reinforcement learning. PLOS Comput. Biol. 18, e1010177 (2022).

32. Rosmalen, R. P. van, Moreno-Paz, S., Duman-Özdamar, Z. E. & Suarez-Diez, M. CFSA: Comparative Flux Sampling Analysis as a Guide for Strain Design. 2023.06.15.545085 Preprint at 10.1101/2023.06.15.545085 (2023).

33. Nogales, J. et al. High-quality genome-scale metabolic modelling of Pseudomonas putida highlights its broad metabolic capabilities. Environ. Microbiol. 22, 255–269 (2020).

34. de Lorenzo, V., Pérez-Pantoja, D. & Nikel, P. I. Pseudomonas putida KT2440: the long journey of a soil-dweller to become a synthetic biology chassis. J. Bacteriol. 206, e00136–24 (2024).

35. Kobak, D. & Berens, P. The art of using t-SNE for single-cell transcriptomics. Nat. Commun. 10, 5416 (2019).

36. Kukurugya, M. A. et al. Multi-omics analysis unravels a segregated metabolic flux network that tunes co-utilization of sugar and aromatic carbons in Pseudomonas putida. J. Biol. Chem. 294, 8464–8479 (2019).

37. Eng, T. et al. Maximizing microbial bioproduction from sustainable carbon sources using iterative systems engineering. Cell Rep. 42, 113087 (2023).

38. Vardon, D. R. et al. Adipic acid production from lignin. Energy Environ. Sci. 8, 617–628 (2015).

39. Tiso, T. et al. The metabolic potential of plastics as biotechnological carbon sources – Review and targets for the future. Metab. Eng. 71, 77–98 (2022).

40. Mokwatlo, S. C. C. et al. Bioprocess development and scale-up for cis,cis-muconic acid production from glucose and xylose by Pseudomonas putida. Green Chem. https://doi.org/10.1039/D4GC03424D (2024) doi:10.1039/D4GC03424D.

41. von Kamp, A. & Klamt, S. Growth-coupled overproduction is feasible for almost all metabolites in five major production organisms. Nat. Commun. 8, 15956 (2017).

42. Tamura, T. MetNetComp: Database for Minimal and Maximal Gene-Deletion Strategies for Growth-Coupled Production of Genome-Scale Metabolic Networks. IEEE/ACM Trans. Comput. Biol. Bioinform. 20, 3748–3758 (2023).

43. Yang, Z. & Tamura, T. DBgDel: Database-Enhanced Gene Deletion Framework for Growth-Coupled Production in Genome-Scale Metabolic Models. Preprint at 10.48550/arXiv.2411.08077 (2024).

44. Santos-Merino, M., Gargantilla-Becerra, Á., de la Cruz, F. & Nogales, J. Highlighting the potential of Synechococcus elongatus PCC 7942 as platform to produce α-linolenic acid through an updated genome-scale metabolic modeling. Front. Microbiol. 14, (2023).

45. Manoli, M.-T. et al. A model-driven approach to upcycling recalcitrant feedstocks in Pseudomonas putida by decoupling PHA production from nutrient limitation. Cell Rep. 43, (2024).

46. Particle swarm optimization. https://ieeexplore.ieee.org/document/488968.

47. Xue, T. & Jieru, Z. Application of Support Vector Machine Based on Particle Swarm Optimization in Classification and Prediction of Heart Disease. in 2022 7th International Conference on Intelligent Computing and Signal Processing (ICSP) 857–860 (2022). doi:10.1109/ICSP54964.2022.9778616.

48. Yang, F., Chen, J. & Liu, Y. Improved and optimized recurrent neural network based on PSO and its application in stock price prediction. Soft Comput. 27, 3461–3476 (2023).

49. Bai, L. et al. Optimizing Metabolite Production with Neighborhood-Based Binary Quantum-Behaved Particle Swarm Optimization and Flux Balance Analysis. J. Comput. Biol. J. Comput. Mol. Cell Biol. 32, 64–88 (2025).

50. Zhang, X.-T., Xu, B., Zhang, W., Zhang, J. & Ji, X. Dynamic Neighborhood-Based Particle Swarm Optimization for Multimodal Problems. Math. Probl. Eng. 2020, 6675996 (2020).

51. Beckham, G. T., Johnson, C. W., Karp, E. M., Salvachúa, D. & Vardon, D. R. Opportunities and challenges in biological lignin valorization. Curr. Opin. Biotechnol. 42, 40–53 (2016).

52. Brienza, F., Cannella, D., Montesdeoca, D., Cybulska, I. & P. Debecker, D. A guide to lignin valorization in biorefineries: traditional, recent, and forthcoming approaches to convert raw lignocellulose into valuable materials and chemicals. RSC Sustain. 2, 37–90 (2024).

53. Ravi, K., García-Hidalgo, J., Gorwa-Grauslund, M. F. & Lidén, G. Conversion of lignin model compounds by Pseudomonas putida KT2440 and isolates from compost. Appl. Microbiol. Biotechnol. 101, 5059–5070 (2017).

54. Salvachúa, D. et al. Metabolic engineering of Pseudomonas putida for increased polyhydroxyalkanoate production from lignin. Microb. Biotechnol. 13, 290–298 (2019).

55. Palmer, R. J. & Staff, U. by. Polyamides, Plastics. in Kirk-Othmer Encyclopedia of Chemical Technology (John Wiley & Sons, Ltd, 2005). doi:10.1002/0471238961.1612011916011213.a01.pub2.

56. van Lent, P., Schmitz, J. & Abeel, T. Simulated Design–Build–Test–Learn Cycles for Consistent Comparison of Machine Learning Methods in Metabolic Engineering. ACS Synth. Biol. https://doi.org/10.1021/acssynbio.3c00186 (2023) doi:10.1021/acssynbio.3c00186.

57. Zhang, J. et al. Combining mechanistic and machine learning models for predictive engineering and optimization of tryptophan metabolism. Nat. Commun. 11, 4880 (2020).

58. Kim, S. et al. PubChem 2025 update. Nucleic Acids Res. 53, D1516–D1525 (2025).

59. Pedregosa, F., et al. Scikit-learn: Machine Learning in Python. J. Mach. Learn. Res. https://inria.hal.science/hal-00650905 (2011).

60. Schneider, P., Bekiaris, P. S., von Kamp, A. & Klamt, S. StrainDesign: a comprehensive Python package for computational design of metabolic networks. Bioinformatics 38, 4981–4983 (2022).

61. Miranda, L. J. PySwarms: a research toolkit for Particle Swarm Optimization in Python. J. Open Source Softw. 3, 433 (2018).

62. Zhao, J. Q., Zhang, C. L., Luo, W. H. & Zhao, J. A Probabilistic Optimal Power Flow Calculation Method with Latin Hypercube Sampling. Adv. Mater. Res. 918, 183–190 (2014).

63. Shields, M. D. & Zhang, J. The generalization of Latin hypercube sampling. Reliab. Eng. Syst. Saf. 148, 96–108 (2016).

64. Bischl, B. et al. Hyperparameter optimization: Foundations, algorithms, best practices, and open challenges. WIREs Data Min. Knowl. Discov. 13, e1484 (2023).

65. SciPy 1.0: fundamental algorithms for scientific computing in Python | Nature Methods. https://www.nature.com/articles/s41592-019-0686-2.

66. Kluyver, T. et al. Jupyter Notebooks – a publishing format for reproducible computational workflows. in Positioning and Power in Academic Publishing: Players, Agents and Agendas 87– 90 (IOS Press, 2016). doi:10.3233/978-1-61499-649-1-87.

67. Lundberg, S. M. & Lee, S.-I. A Unified Approach to Interpreting Model Predictions. in Advances in Neural Information Processing Systems vol. 30 (Curran Associates, Inc., 2017).

68. Escher: A Web Application for Building, Sharing, and Embedding Data-Rich Visualizations of Biological Pathways | PLOS Computational Biology. https://journals.plos.org/ploscompbiol/article?id=10.1371/journal.pcbi.1004321.

